# Spatially aware deep learning reveals tumor heterogeneity patterns that encode distinct kidney cancer states

**DOI:** 10.1101/2023.01.18.524545

**Authors:** Jackson Nyman, Thomas Denize, Ziad Bakouny, Chris Labaki, Breanna M. Titchen, Kevin Bi, Surya Narayanan Hari, Jacob Rosenthal, Nicita Mehta, Bowen Jiang, Bijaya Sharma, Kristen Felt, Renato Umeton, David A. Braun, Scott Rodig, Toni K. Choueiri, Sabina Signoretti, Eliezer M. Van Allen

## Abstract

Clear cell renal cell carcinoma (ccRCC) is molecularly heterogeneous, immune infiltrated, and selectively sensitive to immune checkpoint inhibition (ICI). Established histopathology paradigms like nuclear grade have baseline prognostic relevance for ccRCC, although whether existing or novel histologic features encode additional heterogeneous biological and clinical states in ccRCC is uncertain. Here, we developed spatially aware deep learning models of tumor- and immune-related features to learn representations of ccRCC tumors using diagnostic whole-slide images (WSI) in untreated and treated contexts (n = 1102 patients). We discovered patterns of nuclear grade heterogeneity in WSI not achievable through human pathologist analysis, and these graph-based “microheterogeneity” structures associated with *PBRM1* loss of function, adverse clinical factors, and selective patient response to ICI. Joint computer vision analysis of tumor phenotypes with inferred tumor infiltrating lymphocyte density identified a further subpopulation of highly infiltrated, microheterogeneous tumors responsive to ICI. In paired multiplex immunofluorescence images of ccRCC, microheterogeneity associated with greater PD1 activation in CD8+ lymphocytes and increased tumor-immune interactions. Thus, our work reveals novel spatially interacting tumor-immune structures underlying ccRCC biology that can also inform selective response to ICI.

## Background

Renal cell carcinoma (RCC) is among the 10 most common cancers worldwide and is comprised of several histological subtypes^1^. The clear cell histological subtype (ccRCC) is the most common form of RCC and accounts for the vast majority (75-80%) of metastatic cases^1^. In addition to highly recurrent mutations in hypoxia (*VHL*) and chromatin regulator genes (e.g. *PBRM1, BAP1, SETD2*), ccRCC exhibits extensive genomic intratumoral heterogeneity (ITH)^2^, which was correlated with worse progression free survival in both the TRACERx and TCGA-KIRC cohorts^3–5^. Nuclear grade, an established histopathologic score of tumor nuclei dedifferentiation, is a primary prognostic feature in ccRCC and can provide a histologic description of ITH^6^ to pinpoint cell structures enriched for metastatic potential^7,8^. In addition, high nuclear grade has been associated with increased tumor-infiltrating lymphocytes (TILs) in ccRCC^9^, though whether molecular ITH or its relationship to histologic properties (e.g. grade, TILs) inform immunoresponsive tumor states in ccRCC is uncertain. Indeed, while immune checkpoint inhibitors (ICIs) are a standard therapy in ccRCC, this tumor type defies many conventions about molecular features that associate with selective ICI response identified in other solid tumors^10 11–13^, and both the underlying biology and clinical biomarkers to stratify patients for ICI in ccRCC remain elusive.

Current approaches to simultaneously quantify tumor-intrinsic heterogeneity and its potential relationship to immune microenvironmental interactions in patients are hamstrung by (i) lack of spatial resolution in molecular sequencing, (ii) difficulty with simultaneous multiregional measurements of tumor and immune molecular properties in sufficient cohort sizes, and (iii) practical limitations related to pathologists being incapable of manually perform such measurements from histopathology data at scale. However, by leveraging biologically guided deep learning applied to WSIs, highly detailed evaluation of both established pathology features (e.g. nuclear grade) and novel spatial structures that arise from these features are possible at a scale otherwise intractable via manual pathologist review^14,15^. Thus, we hypothesized that spatially aware deep learning models of ccRCC WSIs could provide a unified understanding of distinct tissue structures that dictate biological and clinical states in ccRCC, and we examined this hypothesis in multiple clinical ccRCC cohorts.

## Results

### Development of a deep learning framework for ccRCC diagnostic images

We first developed prediction models that provide high resolution, quantitative, and human-understandable representations of ccRCC hematoxylin & eosin (H&E) WSIs to identify established pathology features like tumor tissue and nuclear grade at scale^16,17^ (Fig. 1A; Methods, Extended Data Figures 1-8). After quality control analysis, we examined WSIs from 1102 ccRCC patients (n = 421 TCGA-KIRC, 439 CM-025, 208 DFCI-PROFILE, 21 multiblock nephrectomy cases, 13 paired mIF ccRCC cases; Methods). We next trained a second CNN classifier to distinguish low (G2) from high (G4) grade cases in the DFCI-PROFILE cohort, achieving high accuracy at the patient level when evaluated on the TCGA-KIRC dataset using an ensemble of four models (AUROC=0.88; Fig. 1B, Extended Data Fig. 1A). In addition to this binary prediction performance, we also compared the stratification capabilities of continuous nuclear grading to classical pathologist-assigned grades in TCGA-KIRC. We first discretized continuous grade scores into tercile bins to mirror G2/G3/G4 categories, which produced significant patient stratifications for both progression-free interval (PFI) and overall survival (OS) (Fig. 1C, Extended Data Fig. 1B, p <1e-5 [PFI], p < 1e-5 [OS], multivariate log-rank test). Thus, a deep learning computer vision model could both mimic and refine clinically standard categorical nuclear grade assignments in ccRCC.

**Figure 1:**
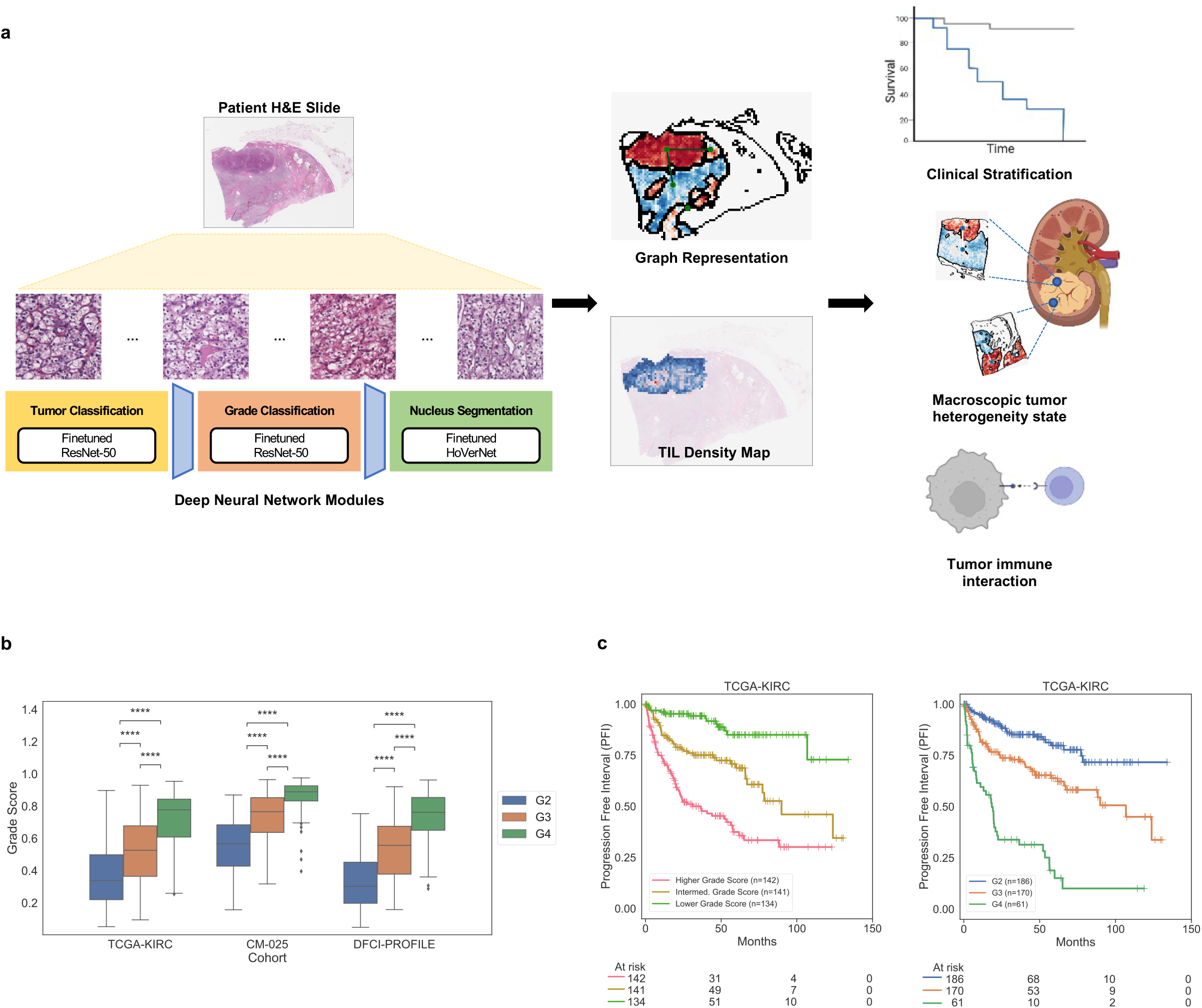
A spatially aware deep learning framework for studying ccRCC. A. Our approach builds a series of biologically relevant prediction models to provide both high resolution and readily human-understandable representations of ccRCC slide images. The first two models identify tumor tissue and grade phenotype within predicted tumor regions, each using a finetuned ResNet-50 convolutional neural network (CNN). A third model identifies tumor infiltrating lymphocytes (TILs) using a finetuned HoVerNet CNN. Local predictions are grouped via watershed segmentation and assembled into graph representations for slide-level description of patients. Computationally inferred patient representations capture both clinically relevant, and biologically informative characteristics of ccRCC. B. Comparison of assigned pathologist grade and grade score on held-out cohorts (TCGA-KIRC, CM-025) in-house training set used for tumor and grade classifier development (DFCI-PROFILE). C. Kaplan-Meier curves for progression free interval (PFI) in TCGA-KIRC based on tercile bins of computationally inferred continuous grade score (left) and assigned pathologist grade (right).

Then, to represent each patient slide compactly, we formed region adjacency graphs (RAGs) that describe where regions of distinct tumor and grade prediction phenotypes occur in a slide, as well as whether these regions directly or indirectly contact one another (Methods). In aggregate, this framework produces a multi-layered, information rich latent representation of ccRCC patient tumor images. Moreover, by condensing the local predictions made by each model, we also represent spatial patterns that arise between these image-derived features.

### Spatial microheterogeneity in ccRCC

Upon inspection of the model representations, we observed a distinct heterogeneity phenomenon in continuous nuclear grade prediction graphs: Some WSIs demonstrated co-occurrence of different grade phenotypes within the same slide, while others were markedly homogeneous. This co-occurrence, which we termed “microheterogeneity”, can be described in two primary (but not mutually exclusive) forms: (i) “proximal”, wherein heterogeneity occurred between tumor tissues that directly contacted one another (Fig. 2A), and (ii) “distal”, wherein stromal barriers or separation in the slide image interrupted the differing tumor tissues (Fig. 2B). We identified microheterogeneity (any proximal or distal occurrences) in 40.6% of TCGA-KIRC cases, and 34.7% of CM-025 (Fig. 2C-D, Extended Data Fig. 9). WSI microheterogeneity was present in varying frequencies within pathologist-assigned grade labels in each cohort, without any consistent pattern between pathologist grade label groups (Fig. 2D, frequency of microheterogeneity = 0.36/0.494/0.317 [G2/G3/G4, TCGA-KIRC], 0.524/0.333/0.230 [G2/G3/G4, CM-025]). To produce a continuous measurement of the amount of microheterogeneity in a single WSI among slides that had microheterogeneity, we calculated the weighted sum of the number of heterogeneous contacts (RAG edges) per WSI, where larger weights are given to contacts with similar tumor region areas (Methods). In two independent ccRCC cohorts, tumors exhibited a wide distribution of microheterogeneity abundance per WSI (Fig. 2E, Extended Data Fig. 10). Thus, in ccRCC WSIs, distinct nuclear grade patterns create microheterogeneity structures that can be quantitatively represented as graphs for further investigation.

**Figure 2:**
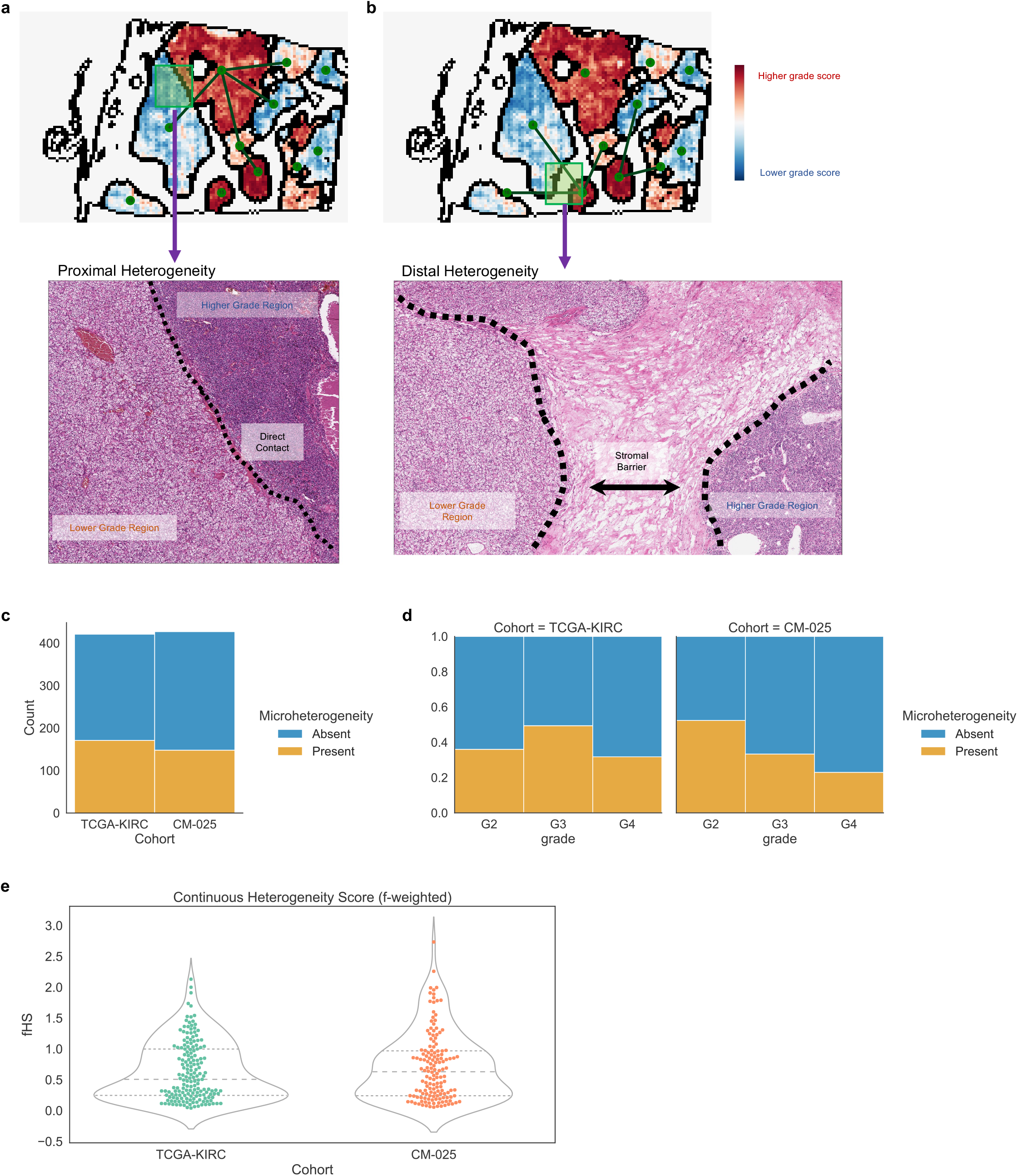
Computationally inferred phenotypic variation in ccRCC. A. Representative example of proximally occurring grade microheterogeneity (dashed line indicating interface of region contact). B. Representative example of distally occurring grade microheterogeneity. C-E: Summary statistics surrounding microheterogeneity in the TCGA-KIRC and CM-025 cohorts. C: Number of patients with/without microheterogeneity. D. Frequency of microheterogeneity by assigned pathologist grade (where available). E. Distribution of continuous heterogeneity score (f-weighted) in non-homogeneous cases, which describes the extent of microheterogeneity in a given slide.

### Establishing the Linkage Between Micro- and Macro-level Heterogeneity

Given the distribution of microheterogeneity abundance per WSI, we then examined how this local, slide-level microheterogeneity related to variation throughout a whole tumor (“macroheterogeneity”). We evaluated a cohort of multiple spatially separated tumor blocks from the same nephrectomy specimen (Fig. 3A-B, Extended Data Fig. 11). For a given patient’s tumor, the maximum microheterogeneity abundance in any single WSI correlated with the presence of microheterogeneity across all WSI from that tumor, and this correlation was not driven by patient sample size (Fig. 3C, Extended Data Fig. 12A-B). In contrast, variation in image-derived grade scores did not correlate with sample size or frequency of microheterogeneity (Fig. 3D; Extended Data Fig. 12C-D). Moreover, subsequent predictive modeling demonstrated that a single WSI could predict microheterogeneity for the remaining WSIs from the same patient (min. log10(Bayes factor) = 3.04; Extended Data Fig. 13; Methods). These findings indicate that observing a single reference slide is predictive of macro-level tumor phenotypes, particularly when that reference slide contains higher grade phenotypes, and that a single ccRCC WSI encodes latent information regarding spatial structures present throughout the tumor.

**Figure 3:**
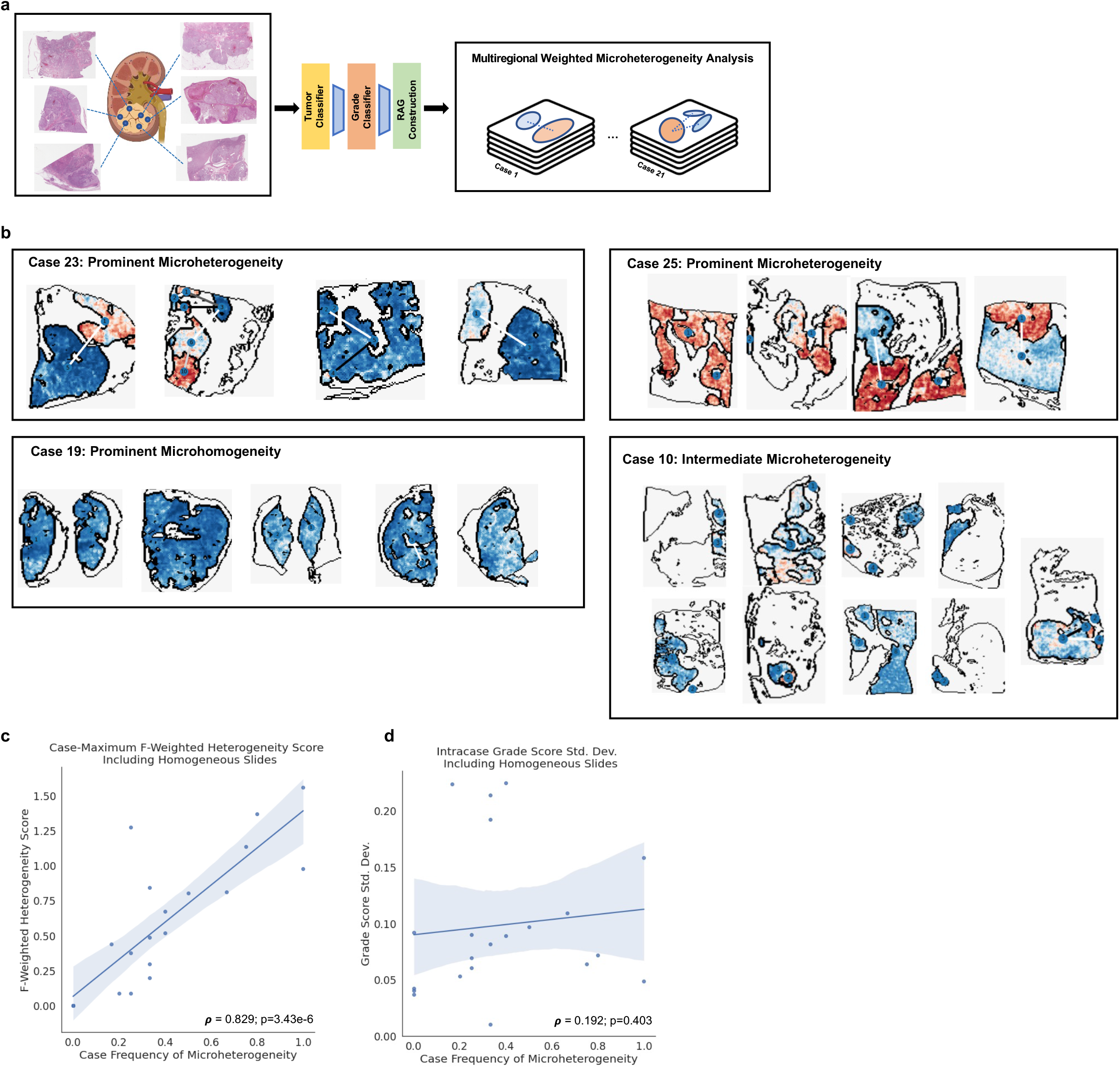
Linkage between microheterogeneiety and whole-tumor variation. A. Schematic for creating a multiregional weighted microheterogeneity analysis using computer vision models. B. Example data collections from four patients, showing RAG plots for the scanned slide of each tissue block. C. Case-wise frequency of microheterogeneity versus the maximum observed f-weighted heterogeneity score, which describes the largest extent of heterogeneity observed in a patient. Statistics aggregated within a given patient’s set of scanned tissue blocks (1 slide per block). D. Case-wise frequency of microheterogeneity versus standard deviation of grade score predictions within the same case. Statistics aggregated within a given patient’s set of scanned tissue blocks (1 slide per block). Pearson’s Rho p-values calculated via exact distribution.

### Molecular correlates of microheterogeneity in ccRCC

Since certain somatic mutations have been associated with macro-level tumor heterogeneity, we subsequently evaluated whether computationally derived microheterogeneity structures from a single WSI were associated with recurrent somatic driver mutations in ccRCC, even though direct prediction of mutations from ccRCC images without multi-layered analysis has thus far been limited^18,19^. WSIs from tumors with somatic *PBRM1* loss of function (LOF), previously associated with molecular ITH, were also associated with a higher frequency of microheterogeneity compared to WSIs from non-LOF tumors (Extended Data Fig. 14). We also examined other common driver mutations in ccRCC and found a similar trend of higher microheterogeneity frequency in *SETD2* LOF mutants, but inconclusive trends for *BAP1* and *PTEN* (Extended Data Fig. 14). Regarding somatic copy number alterations, tumors with 9p21.3 deletions, a molecular feature previously implicated in ccRCC oncogenesis ^20–22^, were enriched for microhomogeneity patterns (Extended Data Fig. 14). Thus, microheterogeneity patterns also encoded features related to recurrent somatic alterations in ccRCC.

### Prognostic relevance of microheterogeneity

Since certain somatic mutations have prognostic value in ccRCC, we assessed whether computationally derived microheterogeneity from WSIs contained additional prognostic information beyond pathologist derived nuclear grade. We compared univariate Cox Proportional Hazards models, using either pathologist assigned grade or computationally inferred continuous grade, to bivariate models that introduced a binary indicator of whether microheterogeneity was observed. In both univariate and bivariate models in TCGA-KIRC, continuous grade had a stronger concordance index (C-Index) for progression free interval (PFI), but not for overall survival (OS) (Extended Data Fig. 15). Within bivariate models for both survival contexts and grading types, the presence of microheterogeneity was negatively correlated with survival (hazard ratios all above 1), most notably in the continuous grade model (Extended Data Fig. 16). Thus, the presence of microheterogeneity in a single localized, untreated ccRCC WSI can identify tumors with poor prognosis and greater metastatic potential, consistent with phenomena previously described through using multi-region molecular profiling^7^.

### Microheterogeneity and response to ICI

In addition to prognostic clinical value, we assessed whether this computer vision derived feature may be predictive for certain ccRCC therapeutics. We assessed spatial microheterogeneity patterns within both treatment arms of CM-025, a phase III randomized clinical trial cohort that compared anti-PD1 blockade (nivolumab) to mTOR inhibition (everolimus) in anti-angiogenic refractory metastatic ccRCC patients (Methods)^13,23^. Presence of microheterogeneity was associated with improved OS and PFS in the ICI arm, but not in the mTOR inhibitor arm (Fig. 4A; Extended Data Fig. 17). Given that continuous grade score correlated with OS for each trial arm, we also examined whether microheterogeneous cases correlated with changes in survival due to having lower overall grade scores. However, within microheterogeneous cases in the ICI arm, grade score did not contribute statistically significant predictive signal for PFS or OS, though it trended toward significance for OS (Fig. 4B, Extended Data Fig. 18). Thus, in CM-025, microheterogeneity was selectively associated with improved response to ICI even though it was a poor prognostic marker in the primary, untreated setting.

**Figure 4:**
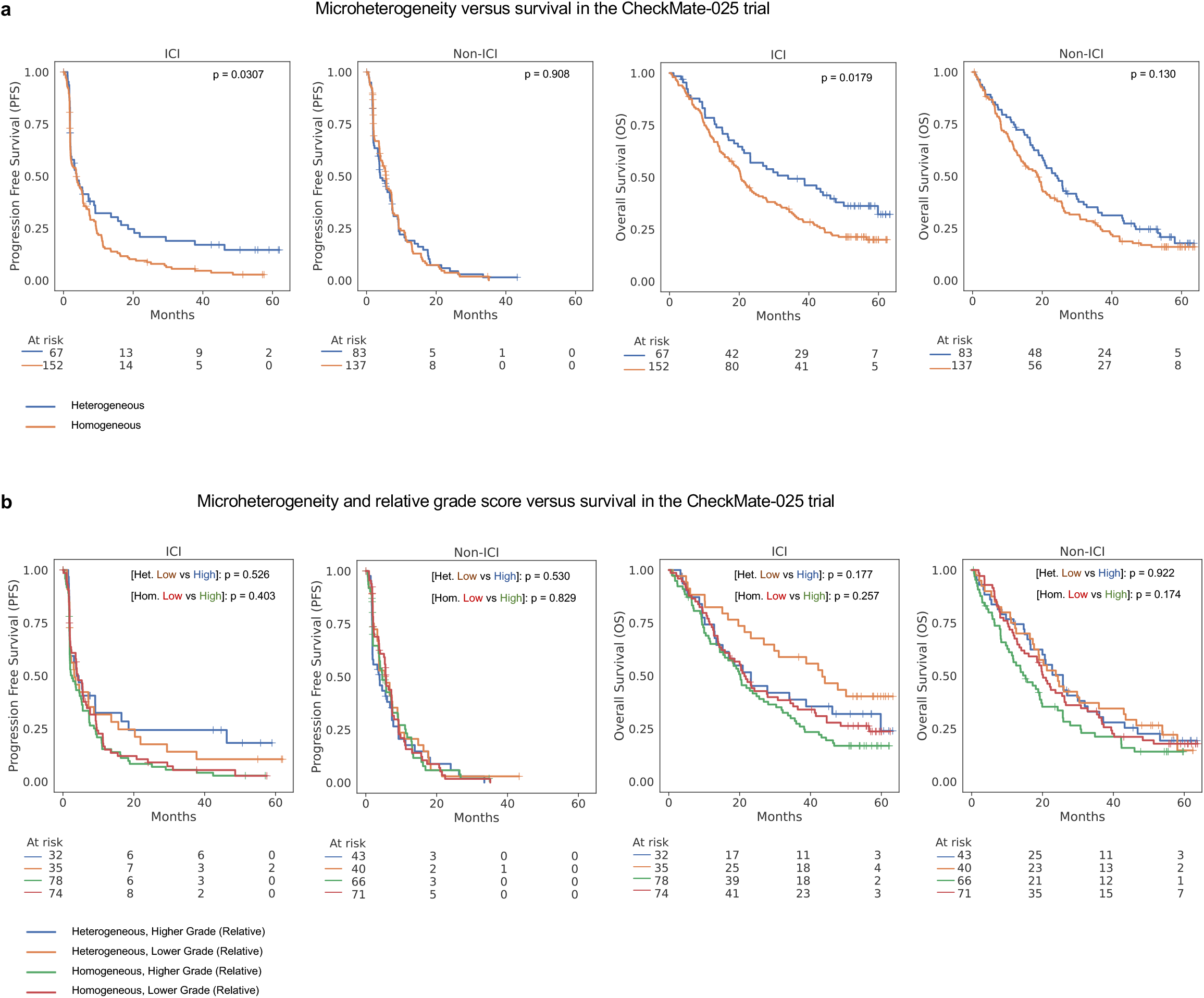
Inferred patterns of grade microheterogeneity associate with improved survival in the CheckMate-025 (CM-025) cohort, but only for immune checkpoint inhibitor (ICI) treated patients. A. Kaplan-Meier curves for overall survival (OS) and progression free survival (PFS) in the CM-025 cohort based on the presence of microheterogeneity. Significance values were calculated via log-rank test. B. Kaplan-Meier curves for overall survival (OS) and progression free survival (PFS) in the CM-025 cohort based on the presence of microheterogeneity, stratifying further based on relative grade score within a grade microheterogeneity category. Significance values were calculated via (pairwise) log-rank test.

### High immune infiltration combined with grade microheterogeneity identifies a further population of ICI responders

Immune infiltration as measured by CD8 immunofluorescence was not associated with response to ICI^13,24^, despite its predictive value in other immune-responsive cancers. We hypothesized that TIL patterns may still be relevant for predicting response to ICI in ccRCC, but joint inference of tumor spatial heterogeneity with TIL patterns are required for adequate context. Thus, we inferred TILs in the CM-025 WSIs and related these features to microheterogeneity (Fig. 5A; Methods). In WSIs with microheterogeneity, highly infiltrated cases associated with improved OS only in the ICI arm (Fig. 5B; p=0.0220, log-rank test). This subset of ICI-treated patients also demonstrated a consistent trend in improved PFS, but did not reach statistical significance (Fig. 5B; p=0.0662, log-rank test).

**Figure 5:**
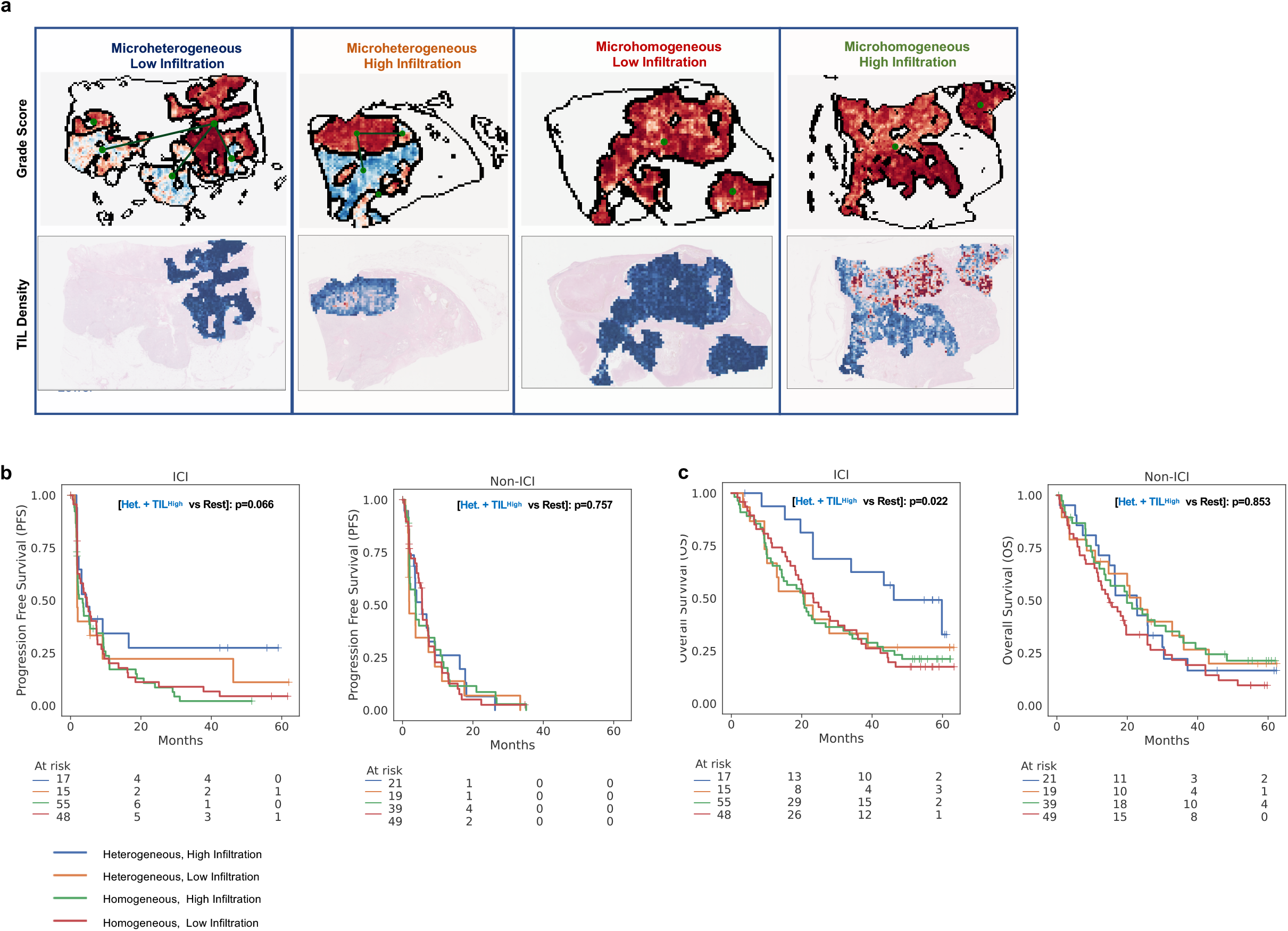
Combining computationally inferred tumor and immune states identifies a further subset of responders to immune checkpoint inhibition (ICI) in the CheckMate-025 trial. A. Representative examples of four classes of patients identified using computational inference, based on the presence of grade microheterogeneity, and the relative abundance of tumor infiltrating lymphocytes in high-grade tumor regions (TIL Density). Top row: representative RAG plots based on grade score (Blue: lower score, Red: higher score). Bottom row: Representative inferred TIL densities (blue: lower infiltration, red: Higher infiltration; uncolored: lower grade regions not considered for TIL density evaluation). B-C: Kaplan-Meier curves for progression free survival (PFS) and overall survival (OS) in both arms of the CM-025 trial based on the groups demonstrated in (a). Significance values were calculated via log-rank test between “Microheterogeneous and High Infiltration” patients and all remaining patients within a trial arm.

We also compared the performance of predictive models that exclusively use image-derived or previously nominated molecular features^13^. For OS in the ICI arm of CM-025, models using computer vision features had similar performance to those only using genomic features (*PBRM1* LOF, 9p21.3 deletion) (Extended Data Figures 19-28; Methods). Moreover, combining these features resulted in net improvements while retaining consistent parameter associations (i.e., *PBRM1* LOF and microheterogeneity each retained positive coefficient weights). We lastly introduced clinical risk covariates into a full parameter model, which produced further improvements to c-index metrics (Extended Data Figures 21, 25; Methods). Taken together, tandem consideration of tumor-intrinsic spatial microheterogeneity and TIL features in WSIs learned by the computer vision models captured meaningful representations of selective ICI response.

### Tumor-immune interactions are more extensive and involve greater CD8+ PD-1 activation in advanced ccRCC

To more precisely understand the tumor-immune spatial interactions identified from WSIs and linked to selective ICI response, we evaluated advanced ccRCC tumors with paired H&E and multiplex immunofluorescence (mIF) images derived from the same tissue (markers = {PAX8, CD8, DAPI, PD1, PDL1, FOXP3})^25,26^. To describe spatial phenotypes, we built a nearest-neighbor graph of CD8+ and tumor cells, and classified cells as “tumor-immune interacting” if they were adjacent to a distinct cell type in the graph (Methods). Through analysis of regions with high tumor-immune interaction density from each patient (Methods), we observed that microheterogeneous tumors had higher CD8+ cell density, while tumor cell density was similar between heterogeneous and homogeneous cases (Extended Fig. Data 31-33). The frequency of tumor cells adjacent to CD8+ cells was higher in heterogeneous cases, suggesting a greater presence of “desert”-like regions of non-infiltrated tumor tissue in homogeneous cases (Fig. 6C, p=0.00215 [Tumor+, tumor-immune], Wilcoxon Rank-Sum test). In contrast, the frequency of CD8+ cells adjacent to tumor cells was similar between heterogeneous and homogeneous cases (Fig. 6C, p=0.418 [CD8+ tumor-immune]). Thus, the observed increase in tumor-immune interaction frequency in microheterogeneous tumors resulted from increased infiltration deeper within tumor-dense regions, rather than a uniform increase across the tumor microenvironment.

**Figure 6:**
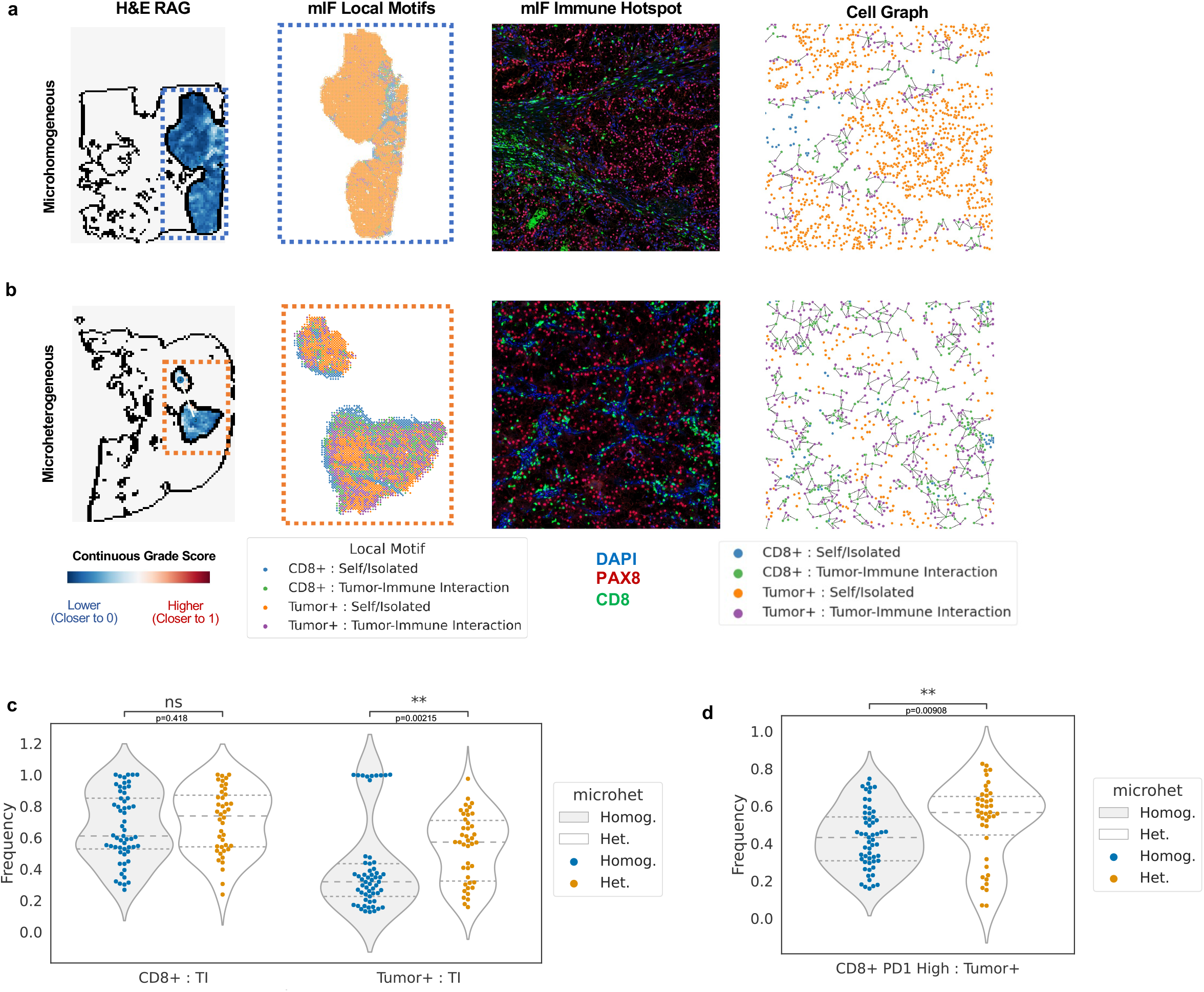
Exploration of microheterogeneity implications in paired multiplexed imaging data. A,B. Representative example of a Microhomogeneous case (A), and Microheterogeneous case (B). H&E Rag: slide-level representation of H&E-inferred tumor and grade properties. mIF Local Motifs: indicates the primary cell type and context present within overlapping 200 pixel windows. mIF Immune Hotspot: representative example of an area of high tumor-immune interaction density. Cell Graph: visualization of tumor and CD8+ cells, and their interaction context; edges are drawn between interacting tumor and CD8+ cells (nearest neighbors in a Delaunay triangulation). C. Comparison of the frequency of tumor-immune interaction within CD8+ cells (left) and tumor cells (right) versus H&E-inferred microheterogeneity status.(TI: “tumor-immune” interaction context). D. Comparison of the frequency of PD1-High within CD8+ cells that interact with tumor cells versus H&E-inferred microheterogeneity status. Significance calculated via Wilcoxon rank sum test. p-value annotation legend: ns: p <= 1.00e+00 *: 1.00e-02 < p <= 5.00e-02 **: 1.00e-03 < p <= 1.00e-02 ***: 1.00e-04 < p <= 1.00e-03 ****: p <= 1.00e-04

We lastly asked whether any of these observed differences related to tumor-immune cell subtypes, specifically PD1 low/high CD8+ and PDL1 low/high tumor cells. In general, PD1 high CD8+ cells were common, and PDL1 high tumor cells were sparser (PD1 median freq. = 0.480, PDL1 median freq. = 0.150; Extended Data Fig. 33). Within CD8+ cells engaged in a tumor interaction, microheterogeneous cases had a higher frequency of PD1 high cells compared to homogeneous cases (Fig. 6D, p = 0.00908, Wilcoxon Rank-Sum test). Thus, spatial microheterogeneity structures in ccRCC, which exhibited enrichment for ICI response, may foster an immune compartment that is both more tumor experienced and abundant.

## Discussion

Simultaneous quantitative measurements of key tumor and microenvironmental properties that represent distinct modes of oncogenesis, evolution, and immune evasion may unlock new insights in ccRCC biology and potential modes of patient stratification. To this end, we developed a series of biologically informed neural network models to perform spatially aware computer vision analysis on multiple independent ccRCC cohorts. In doing so, we produced a continuous, quantitative, and automated grading approach that reproduces existing manual histological assessments of nuclear grade and provides comparable prognostic value without interobserver variability. More importantly, by formalizing tumor phenotype predictions into spatial maps and subsequent region adjacency graphs using a single WSI per patient, we discovered histological intratumoral heterogeneity properties not feasibly measurable by manual review that were informative for multiple phenomena, and represented patterns present throughout a patient’s tumor. Namely, the graph-based microheterogeneity feature contained additional prognostic value beyond established pathology scores, as well as predictive value specifically for response to ICI in CM-025. Furthermore, this feature correlated with a series of molecular characteristics, such as *PBRM1* LOF, and thus may provide a unified histological representation for connecting clinically relevant molecular features. Upon simultaneously integrating tumor and immune microenvironmental features, we identified a subset of ICI responders enriched for microheterogeneity and a higher degree of TILs. Moreover, microheterogeneity in advanced ccRCC associated with greater PD1 activation in CD8+ lymphocytes and a greater extent of tumor-immune interaction, suggesting a more active tumor-immune interaction landscape that is more likely to respond to ICI. Taken together, these findings suggest that tumor and immune features of ccRCC can be jointly considered in a spatially aware manner to guide biological and clinical investigations using widely available H&E WSIs.

There are several challenges and limitations to this analysis. The histological data analyzed from CM-025 consisted of pre-treatment primary tumor samples, and thus may differ from the tumor state at the time of trial accrual due to ongoing tumor evolution. As such, the specimens we analyzed may be uncoupled from eventual metastatic progression. Similarly, larger sample sizes in additional clinical cohorts are necessary to generalize these findings to the evolving combination treatment landscapes of ccRCC, and additional histologic features could be added to our model framework (e.g. necrosis, TIL subtypes). While we were able to provide an orthogonal glimpse at the specific cell populations that might underlie the tumor-immune phenotypes associated with microheterogeneity, our analysis of paired mIF and H&E data also had key limitations. In particular, the sample size was small and composed of varying biopsy sites, and larger paired cohorts representing diverse biopsy sites will guide extensions of these observations.

Taken together, we propose spatially aware deep learning models that build upon inference of known histological features (e.g. nuclear grade) to learn interacting new features (graph-based microheterogeneity) and reveal distinct oncogenic paths and ICI response phenotypes in ccRCC. The occurrence of microheterogeneity and its predictive capacity for PFS and OS in ICI warrants further study, including via model systems, to unravel how this phenomenon influences tumor evolution and anti-tumor immunity. Broadly, the use of biologically guided computer vision strategies for cancer histopathology to automatically infer tumor and microenvironmental features, their respective higher-order interactions, and their relationship to molecular and clinical states may have general utility across tumor types and therapeutic modalities.

## Methods

### Clinical Cohorts

Three distinct patient cohorts were used in the analysis: TCGA-KIRC, the CheckMate 025 (CM-025) phase III clinical trial (NCT01668784), and the DFCI-PROFILE of ccRCC patients from the Dana-Farber Cancer Institute (also under DFCI IRB #20-293 and 20-376). Additional datasets include a set of multi-block nephrectomy samples from Dana-Farber/Brigham (under DFCI IRB #20-293 and 20-376), and multiplexed immunofluorescence data from Dana-Farber/Brigham the ImmunoProfile project (under DFCI IRB #20-293 and 20-376).

### Data Acquisition

For the TCGA-KIRC cohort, we obtained clinical data, and normalized bulk RNA and genomic sequencing from the GDC PanCanAtlas (https://gdc.cancer.gov/about-data/publications/pancanatlas), and downloaded whole-slide H&E stained diagnostic images from the ISB-CGC mirror of TCGA data. For the CheckMate 025 cohort, we directly obtained H&E stained diagnostic images from the Signoretti Lab via an established Bristol Myers Squibb IION agreement, and used clinical and molecular data previously generated by Braun et al., 2020 (European Genome-Phenome Archive: EGAS00001004290, EGAS00001004291, EGAS00001004292). DFCI-PROFILE images were obtained via the Dana-Farber/Brigham and Women’s PROFILE project. Multiplexed Immunofluorescence (and accompanying H&E images) were obtained via the Dana-Farber/Brigham and Women’s ImmunoProfile project. Multi-block nephrectomy images were obtained from the Signoretti Lab. These images are available upon request with provision of IRB and adherence to institutional policies regarding storage security and other parameters.

### Truncating mutation categorization

When considering somatic alterations, we consider mutations only if they are truncating (likely loss of function). Within MAF annotation data, this comprised the following variant categories: {‘Nonsense_Mutation’, ‘Frame_Shift_Ins’, ‘Frame_Shift_Del’, ‘Splice_Site’}.

### Image Quality Control

Quality control of H&E whole-slide images was performed using the HistoQC^27^ toolkit. A full set of modules used is available as a ‘.ini’ file. Custom examples used to train pen detection modules are available in a forked repository (https://github.com/jmnyman/HistoQC)]. Following HistoQC filtration, we further removed small slide images with fewer than 500 tiles (512px at 20X resolution).

### Cross Validation

Training and validation were performed in the DFCI-PROFILE cohort, with additional testing done in the TCGA-KIRC and CM-025 cohorts. In subsequent finetuning experiments for tumor and grade classification, we utilized 4-fold cross-validation. Each dataset was split into folds at the patient level to ensure no patient-level information bled between training and validation contexts. Datasets were then subsampled to ensure a balanced label composition. Following balanced patient-level fold creation, we then sample a fixed number (per specified hyperparameters) of 512 pixel tiles (20X) from each patient slide to create folds that were label-balanced.

### Tumor Classification

A model to classify general renal cell carcinoma tumor tissue versus adjacent normal stromal tissue was trained using pixel-level pathologist annotations from the DFCI-PROFILE cohort (n=36 slides). Following quality control, training data slides were split into 512 pixel tiles (20X), and assigned labels according to pathologist annotations (ie, whether a region contained tumorous tissue or not). A pretrained ResNet-50 neural network model was then finetuned using color jitter, and the highest performing model in a series of 4-fold cross validation was selected for subsequent inference. All neural network architecture and training code used the PyTorch and PyTorch-Lightning libraries, and 1-2 NVIDIA Tesla V100 GPU units on Google Cloud VM instances ^28,29^. Hyperparameters used for training are available in the project repository.

### Tumor Grading

A second finetuned ResNet-50 neural network model was trained to distinguish low (G2) from high (G4) ccRCC cases from the DFCI-PROFILE cohort (n=190 slides). This cohort contained samples collected prior to and following the adoption of the WHO/ISUP grading changes, and as such contains both Fuhrman and WHO/ISUP grades ^6^. These conventions share significant overlap and are generally highly concordant ^30^. Additionally, manual review for sarcomatoid and rhabdoid (S/R) tumor content was previously performed, and cases with S/R content were upgraded to G4 if previously assigned a lower grade to ensure greater concordance with WHO/ISUP guidelines. Following quality control, training data slides were again split into 512 pixel tiles (20X), and tiles were assigned labels according to pathologist annotation of the source slide. Tiles were only considered for model training if their predicted probability of containing tumor tissue was >= 0.7, and slides were restricted to those with at least 500 putative tumor-containing tiles. A pretrained ResNet-50 neural network model was again finetuned using heavy color jitter. An ensemble composed of each model trained in 4-fold cross validation was then used for subsequent inference, taking the average across all model softmax outputs to make predictions. All neural network architecture and training code used the PyTorch and PyTorch-Lightning libraries, and 1-2 NVIDIA Tesla V100 GPU units on Google Cloud VM instances ^28,29^. Hyperparameters used for training are available in the project repository.

### Inference Post-Processing

Following tumor and grade inference, tile-level model scores were smoothed using uniform nearest neighbor averaging (n=4 nearest tiles). For grade score smoothing, we considered tiles if their predicted probability of containing tumor tissue was >= 0.5. Smoothed tumor and grade scores were used for all downstream analysis.

### Phenotype Segmentation

Regions of tumor tissue were identified using watershed segmentation in scikit-image ^31^. We first performed segmentation on smoothed tumor prediction scores, and classified regions as “tumor” if their average segment score was >= 0.7. Following an initial watershed segmentation, regions were merged if they were similar (region score difference < 0.2). These putative tumor regions were then considered for secondary segmentation using smoothed grade scores to identify regions of distinct grade.

Furthermore, a slide-average grade score was obtained using the average grade score across all putative tumor area.

### Adjacency Descriptions

Following watershed segmentation of tumor and grade scores, we represented each slide as a region adjacency graph^32^ (RAG) to describe the connectivity of each region produced, wherein directly contacting regions are assigned an edge in the graph. Small area nodes were removed (n < 50 total tiles) following RAG construction. Subsequently, we performed a series of segmentation expansions to recover missing connectivity locally (e.g., regions that are visually in contact, but are separated by a thin layer of non-tumor predicted area), and also to describe long-range differences (e.g., regions that are distinct in grade score, but separated by 10+ tiles). RAG edges forming either directly or at an expansion distance of 1 tile were classified as “proximal”, and those forming at an expansion distance of up to 25 tiles were classified as “distal”. In analyses using TIL predictions, we only considered edges containing at least one node with an average grade score above 0.8 (see *Tumor Infiltration Classification*).

### Heterogeneity Description

Patients with at least one proximal or distal RAG edge were classified as “heterogeneous”, and those lacking any edges as “homogeneous”. In analyses using TIL predictions, we only considered proximal/distal edges containing at least one node with an average grade score above 0.8 (see *Tumor Infiltration Classification*). To describe microheterogeneity continuously, we derived two related metrics. First was a “total-weighted” heterogeneity score (t-HS), we calculated a weighted sum of the number of RAG edges, wherein each edge was weighted by the total fractional tumor area occupied by the node pair involved in that edge (e.g., if two nodes comprise nearly all of the tumor area, that edge is highly weighted, while smaller regions contribute less to the sum). The second score was “f-weighted” (f-HS), and instead used the harmonic mean of the area fractions of each node in an edge to produce a weight, which describes both the contribution scale and balance of the nodes involved in that edge.

### Reference Slide Modeling

Null models were configured for each case based on the number of blocks available for a given patient (1 H&E stained slide per block, n=21 patients, minimum 3 blocks, average = 4.57 blocks, median = 4 blocks, max = 10 blocks), and a Beta prior was set according to the empirical observations of microheterogeneity in the CM-025 cohort (prior parameters: a=148, b=279). Alternative models for each patient were configured by first setting a uniform prior (a=1, b=1), and then updating based on the microheterogeneity status of a reference slide (e.g., {a=2, b=1} when observing microheterogeneous reference). We selected the reference slide based on grade score in 4 different ways: highest slide-average, highest by segment, highest by segment (with 10% area minimum), and highest by segment (with 25% area minimum). We also considered near-ties, considering a tie if two candidate grade scores were within 0.01 of each other, taking the average log likelihood of the competing alternative models for a given patient when comparing to the null in downstream testing. Comparison testing was done with a log-likelihood ratio test. Bayes factors were calculated analytically under beta-binomial distributions.

### Tumor Infiltrating Lymphocyte Inference

A HoVerNet model trained on the PanNuke dataset was used for nuclei segmentation ^17,33^. We leveraged the implementation and pretrained model from the PathML toolkit ^34^ for computational pathology (https://github.com/Dana-Farber-AIOS/pathml). While this pretrained model produces accurate nucleus segmentation, its subtype classification notably fails on clear cell renal carcinoma, likely due to a near-absence of this histology within PanNuke. Consequently, we finetuned the classification head of the model to predict tumor-context vs stromal-context nuclei using a pseudo-labeling scheme; nuclei in a tile were randomly assigned “tumor nuclei” labels proportionally to the predicted probability produced by the tumor classifier (ex., 90% of nuclei randomly assigned “tumor nuclei” if tumor score == 0.9). Following inference, we further stratified nuclei predicted to be “tumor-context nuclei” to distinguish tumor cells from infiltrating lymphocytes (TILs) using heuristic cutoffs chosen via manual pathologist review; lymphocytes were selected by a combined criteria of increased circularity, smaller area, and darker pixels.

### Tumor Infiltration Classification

Following nuclei inference, we aggregated nuclei calls at the tile-level, and classified a tile as “infiltrated” if it contained 14 or more TILs, a cutoff selected by maximizing concordance with pathologist annotations for the presence of lymphocytes in a given tile (Extended Data Fig. 4; bootstrapped AUROC comparing “infiltrated” vs “non-infiltrated” tile-level labels). We then considered region-level descriptions of infiltration, describing the proportion of tiles above the “infiltrated” cutoff as the “area infiltration fraction”. Next, we binarized samples into “low” versus “high” infiltration by splitting at the median area infiltration fraction value (cutoff = 15.16%). This was restricted to high-grade regions (grade score >= 0.8) to avoid excessive false positive infiltration calls, as lower-grade ccRCC nuclei can be visually ambiguous from TILs, even to expert pathologists (n=256 patient slides post filtration) (See Extended Data Figures 5-7). Area infiltration fraction as determined by H&E-inference showed general agreement with CD8+ immunofluorescence measurements where overlap existed (See Extended Data Fig. 7]).

### Survival Analysis

Survival analysis in the TCGA-KIRC And CM-025 cohorts was performed using the python package *Lifelines* ^35^. Kaplan-Meier regression and plotting was performed using the *KaplanMeierFitter* function with default parameters, and multivariate Cox Proportional Hazards regression was performed using the *CoxPHFitter* function with moderate regularization (L1 ratio = 0.1 [multivariate models], L1 ratio = 0 [univariate models], penalizer scale = 0.1 [multivariate models], penalizer = 0.0 [univariate models]). We also further excluded slide images with fewer than 200 tiles predicted to contain tumor tissue. We considered only slides obtained from primary biopsy sites, excluding metastatic biopsies to remain consistent between cohorts. When annotations were available, we excluded Grade 1 (G1) cases due to their rarity. We only considered cases where watershed segmentation successfully produced at least one segment containing 50 or more tiles with an average tumor score >= 0.7. When describing TIL infiltration content in CM-025 in Kaplan-Meier curves, we used binary (lower/higher) groups as described above, and continuous area infiltration fraction for Cox modeling.

### Image Registration

We adapted PathFlow MixMatch, displacing an input H&E image against a fixed mIF image at 1.25X, and using GPU acceleration (Tesla V100) when learning each case’s alignment/displacement tensor. Learned displacement tensors were then used to shift H&E-based grade segmentation maps into the same coordinate space as mIF data. These aligned maps were then used in K-nearest neighbors regression to assign cell predictions in the mIF data to a grade segmentation label. Since the image pairs are not from the same exact tissue section, alignments were assessed visually via overlay to assess quality, resulting in 13 total passing cases. Within successful alignments, putative tumor regions were manually reviewed, resulting in omission of two false-positive regions (adjacent metastatic tissue misclassified as “tumor”).

### Multiplexed Immunofluorescence Image Preprocessing

To first predict cellular locations, we used a pretrained Mesmer model ^36^, which produced a candidate mask of cell segmentations for each mIF WSI. We used DAPI as the “nuclear channel”, and PAX8 with CD8 as the “membrane channels”. Full resolution (20X) mIF images were broken into 10,000 pixel bands as batch inputs to Mesmer using GPU acceleration (Tesla V100). A subset of images (3) that failed this batch procedure were re-run with 2500 pixel square tiles as batched inputs. To quantify area-normalized cellular expression, we used the Ark analysis toolkit’s ‘create_marker_count_matrix’ function ^36,37^, which produces a description of each predicted cell segmentation that contains both morphological and arcsinh-transformed, area-normalized expression values.

### Cell Phenotype Calling

Following expression quantification, we inspected the histograms of each case’s channel values, as well as the ratio of CD8+ : Autofluorescence, and determined manual cutoffs to gate each primary cell population (CD8+ vs Tumor+ vs ungated). These cutoffs were then used to make a coarse-grained estimate of each cell subpopulation. We then performed an orthogonal clustering analysis, using cell expression and morphology features produced by the Ark toolkit (*{centroid_dif, num_concavities, convex_hull_resid, major_axis_equiv_diam_ratio, perim_square_over_area, arcsinh(Cell Area)}*) and the Louvain method for community detection in scanpy (number of principal components = 5, nearest neighbors=15, cluster resolution=10) ^38,39^. Cells with low DAPI (<7 arcsinh units) were also excluded. Clusters with outlier morphology (>50% of its cells having 3+ features outside of 5th/95th percentile values), or high autofluorescence-to-CD8+ signal (>35% below case-specific CD8+ : Autofluorescence ratio cutoffs) were excluded. Remaining clusters were assigned to “CD8+” or “Tumor+” identities if at least 60% of a cluster’s cells were assigned that label when using purely manual cutoffs. Remaining cells were labeled “ungated” and excluded from downstream analysis. To determine cell subpopulations, we subsetted each primary cell population (CD8+, Tumor+), and fit a linear regression model of autofluorescence vs submarker expression, using the resulting residuals as a noise-corrected expression value. Resulting submarker distributions (PDL1 for Tumor+ cells and PD1 for CD8+ cells) were inspected, and binary cutoffs were chosen for each case individually. We lastly performed a filtering of false-positive tumor cell predictions which exhibited high PAX8 and high DAPI (likely to be B cell lineage), and again used manual histogram inspection to remove the DAPI-high subpopulation.

### Cell graph construction

To construct a graph of cell-interactions, we first removed ungated cells, and then performed Delaunay triangulation with a maximum radius of 100px to form a parsimonious nearest neighbor graph. We then defined “self” interactions as edges between cells of the same type (e.g. CD8+), and “tumor-immune” interactions as those occurring between CD8+ and Tumor+ cells. Cells disconnected from the graph were deemed “isolated”, and grouped with “self” interactions for downstream analysis.

### Immune Hotspot Analysis

To select regions of interest with high tumor-immune activity, we split full resolution mIF WSI data into tiles 2000px (approx 1mm) wide, and further selected for regions with at least 50% area overlap with H&E-inferred tumor region predictions. We then filtered for regions with at least 50 CD8+ and 50 Tumor+ cells that were engaged in tumor-immune interactions. From each patient slide, we sampled up to 10 hotspots, selecting those with the most CD8+ density involved in tumor-immune interactions (min=2 samples, mean=9.0 samples; 42 total microheterogeneous samples [n=5 slides], 57 total microhomogeneous samples [n=6 slides]) (See Extended Data Figures 28-29). Two patients lacked any hotspots and were excluded from this analysis.

### Statistical Testing

All statistical analysis was performed using python 3. For comparison of group counts, Fisher’s Exact test was used via the scipy function *fisher_exact* ^*40*^. Other continuous, score-based comparisons were performed using a two-sided Wilcoxon rank-sum (Mann-Whitney U) test using the *statannotations* package to directly annotate Seaborn plots with p-value results ^41,42^. For survival analyses, cohort subgroup survival distributions were compared using the log-rank test using the *multivariate_logrank_test* and *pairwise_logrank_test* functions in *Lifelines*. For Cox models, the concordance index (C-Index) and (Log) Likelihood Ratio Test (LLRT) were used to evaluate goodness of fit. When comparing continuous to categorical grade, the relative likelihood was estimated using the partial AIC produced for each Cox model, and interpreted as the probability one model minimizes the AIC of the other. Barplot error bars indicated standard error. Boxplot elements are as follows: center line, median; box limits, upper and lower quartiles; whiskers, 1.5 interquartile range (IQR); points, outliers past 1.5 IQR. Violinplot dotted interior lines indicate median, and upper and lower quartiles.

## Data Availability

Restrictions apply to the availability of the raw in-house and external data, which were used with institutional permission through IRB approval for the current study, and are thus not publicly available. Please email all requests for academic use of raw and processed data to DFCI Contracts Team. All requests will be evaluated based on institutional and departmental policies to determine whether the data requested is subject to intellectual property or patient privacy obligations. Data can only be shared for non-commercial academic purposes and will require a formal Data Use Agreement.

## Code Availability

Code used to perform the analyses described in this study will be made available in a public github repository upon publication.

## Acknowledgements

This work was supported in part by Bristol Myers Squibb through its International Immuno-Oncology Network.

The results shown here are also in part based upon data generated by the TCGA Research Network: https://www.cancer.gov/tcga. This work was supported by Dana-Farber/Harvard Cancer Center Kidney SPORE (P50CA101942-15). This work was also supported in part by the National Institutes of Health (R37 CA222574, R01CA227388, P50CA191842 [E.M.V.A.]; T32GM007753 [N.M.]; P30CA016359 [D.A.B.]; F31 CA250136 [J.N.]), a Dunkin’ Donuts Breakthrough Grant [J.N., E.M.V.A.], and the DOD Academy of Kidney Cancer Investigators (KC190128) [D.A.B.]. T.K.C. is supported in part by the Kohlberg Chair at Harvard Medical School and the Trust Family, Michael Brigham, and Loker Pinard Funds for Kidney Cancer Research at DFCI.

## Author Contributions

Conceptualization - J.N., E.M.V.A, S.S., D.A.B., S.S., T.K.C.; Formal analysis - J.N. ; Interpretation of the data - S.N.H, B.J., N.M., C.L., Z.B., T.D., D.A.B., K.B., B.M.T; Resources - R.U., J.R.; Data curation & annotation - T.D., Z.B., C.L., S.S., S.R., K.F., B.S.; Writing (original draft) - J.N., E.M.V.A; Writing (reviewing and editing) - S.S., D.A.B., S.S., T.K.C., B.J., N.M., S.N.H, K.B., B.M.T; Visualization - J.N.; Supervision - E.M.V.A, S.S., T.K.C.; Funding acquisition - E.M.V.A, S.S., T.K.C.

## Extended Data

**Extended Data Figure 1:**
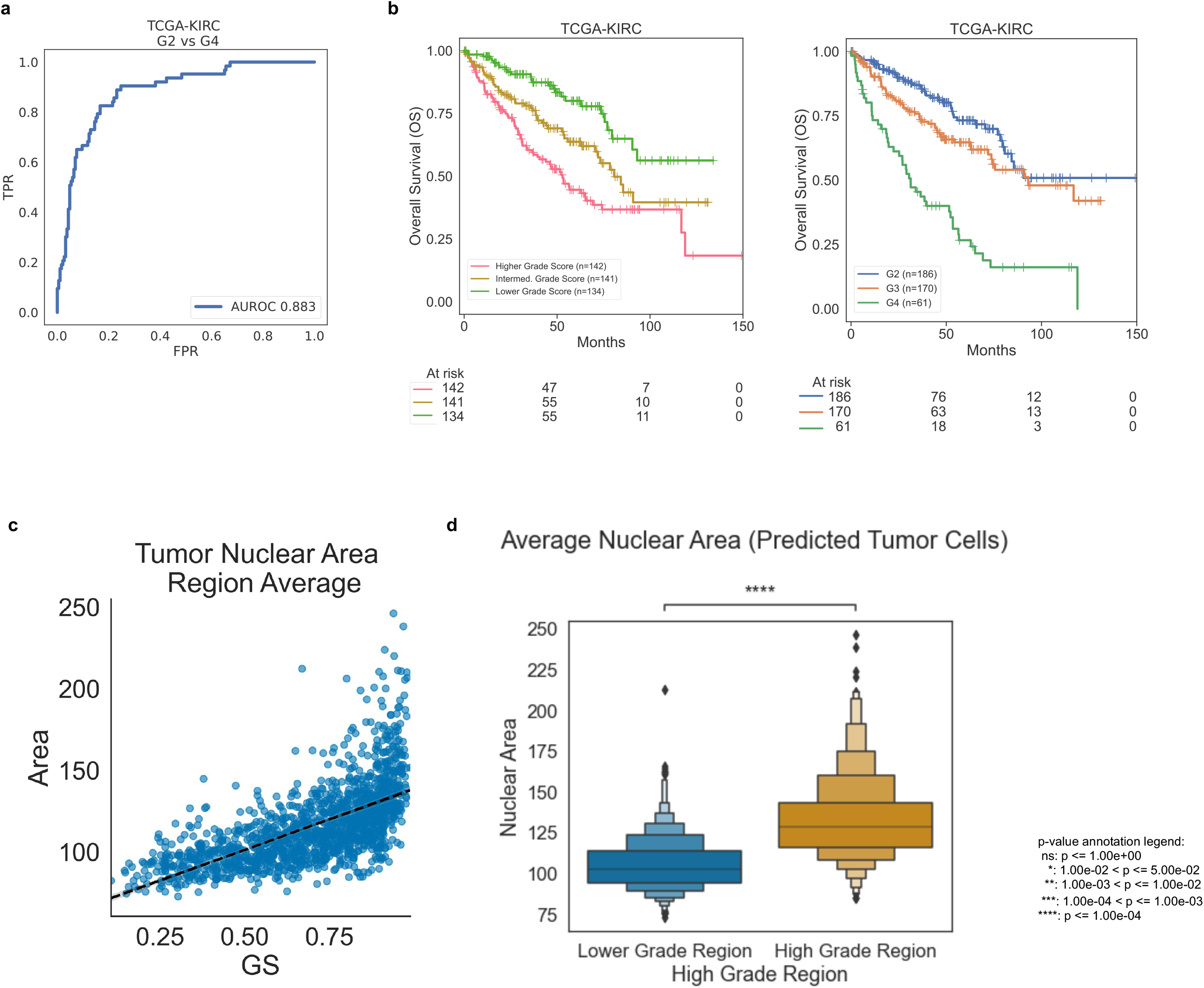
Evaluation of grade neural network model. A. Receiver operator characteristic curve (ROC) for evaluating performance of a grade classifier on TCGA-KIRC (*AUROC: area under ROC curve statistic; TPR: true positive rate; FPR: false positive rate*). B Kaplan-Meier curves for overall survival (OS) in TCGA-KIRC based on tercile bins of computationally inferred continuous grade score (left) and assigned pathologist grade (right). C. The average area of predicted tumor nuclei versus grade score (GS), aggregated over distinct tumor regions per WSI. D. Dichotomizing regions based on high grade designation (average score above 0.8).

**Extended Data Fig. 2:**
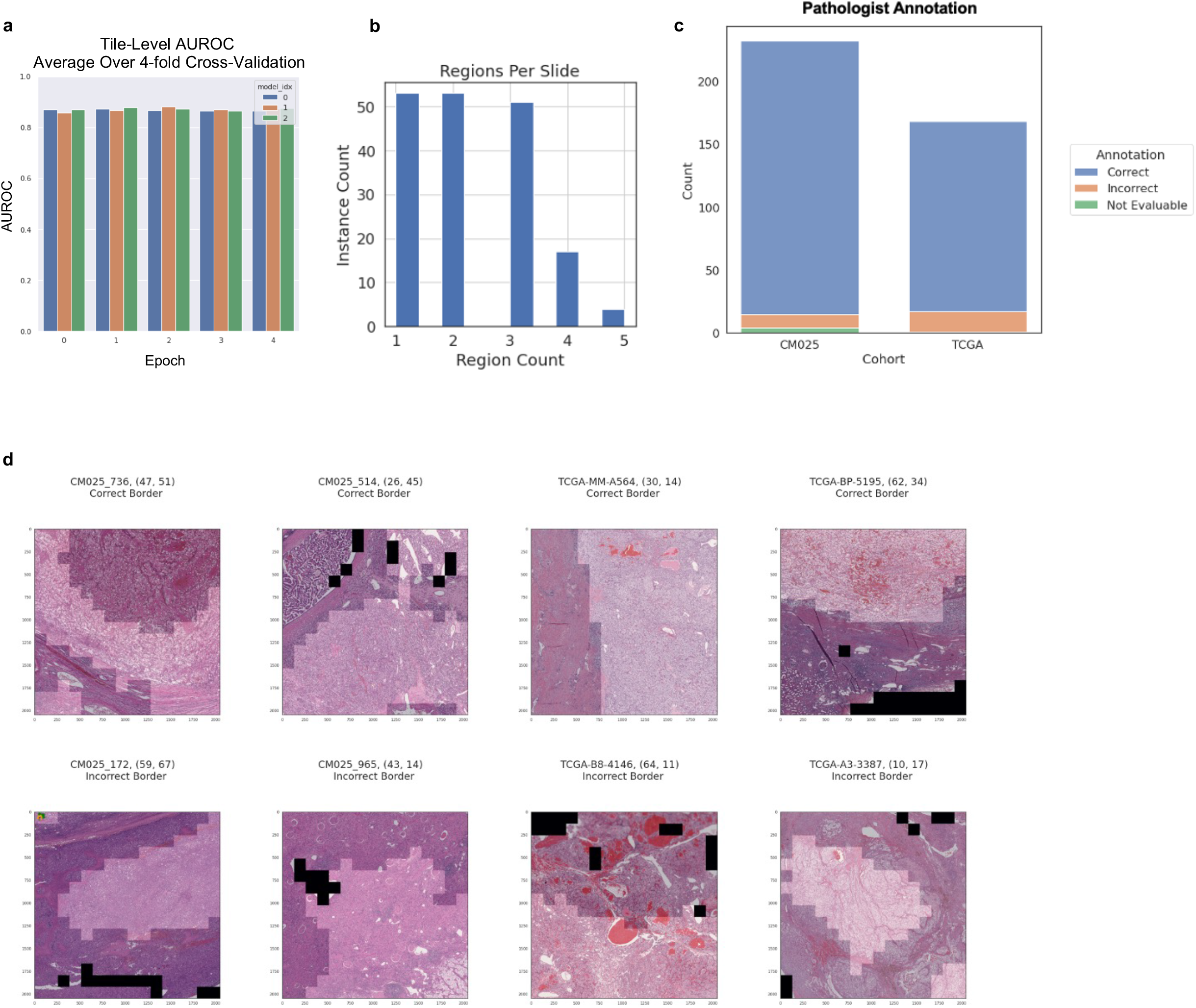
Evaluation of tumor region segmentations by pathologist. A. Tile-level AUROC for tumor vs non-tumor prediction, averaged over 4-fold cross-validation. (Model 0: Finetuned ResNet-18, Model 1: Finetuned ResNet-50 (Selected for downstream), Model 2: Modified ResNet-50.) B. Number of regions examined per sampled slide. C. Distribution of pathologist assessments by cohort. D. Examples of correct and incorrect tumor-stroma borders.

**Extended Data Figure 3:**
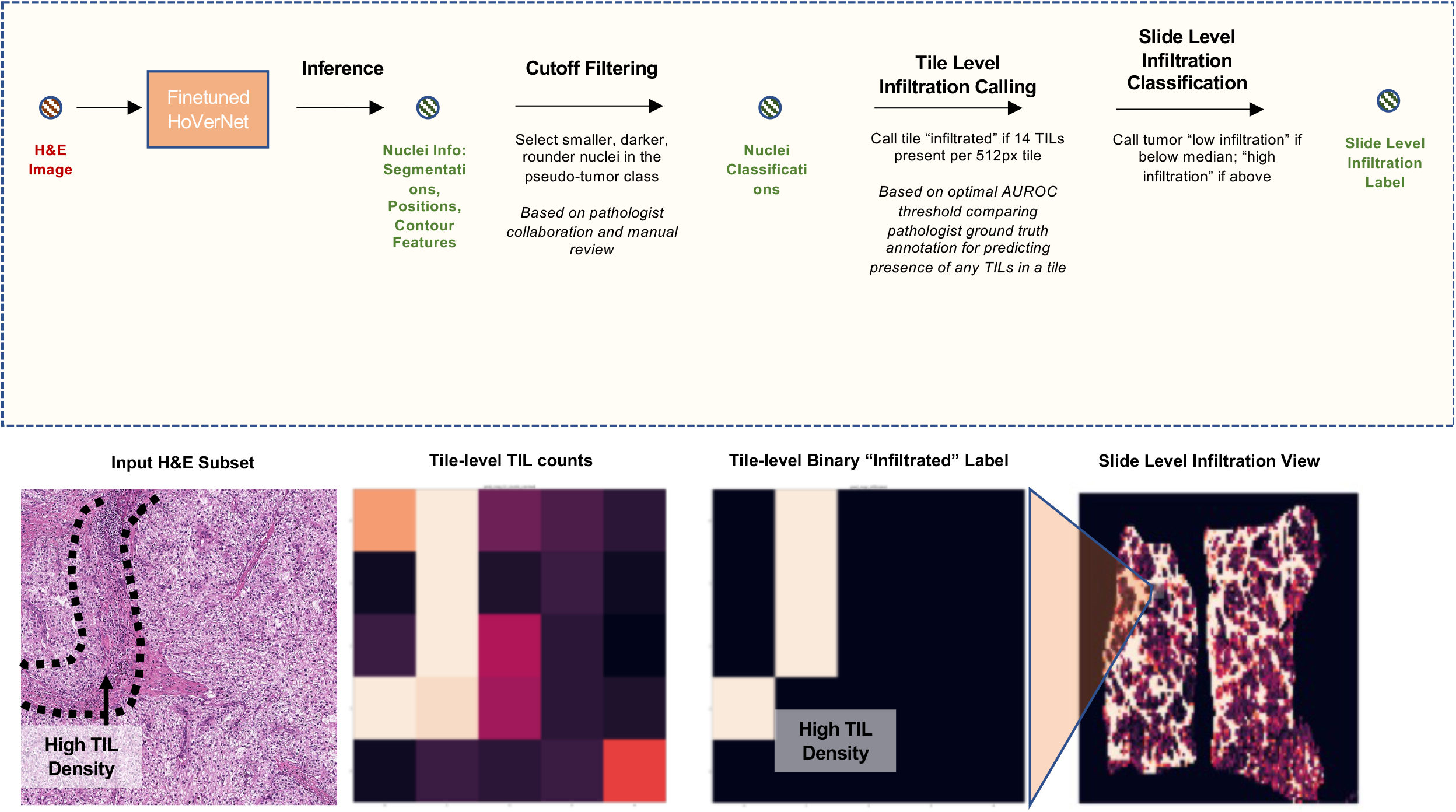
TIL inference process description. Top: sequential descriptions of inference process. Bottom: visual example for illustration.

**Extended Data Figure 4:**
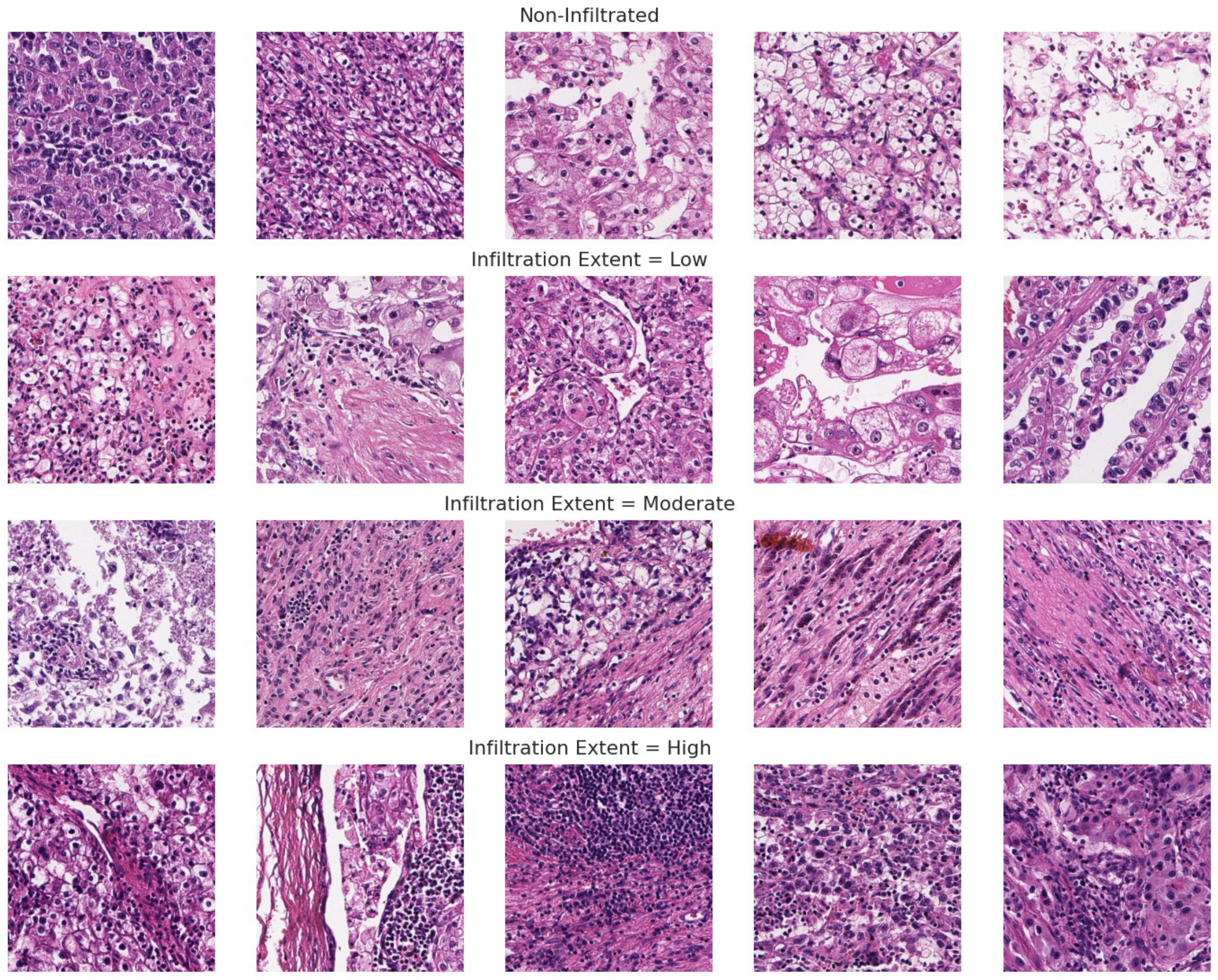
Ground truth/pathologist-labeled examples for TIL infiltration extent used for evaluating tile level TIL thresholds.

**Extended Data Fig. 5:**
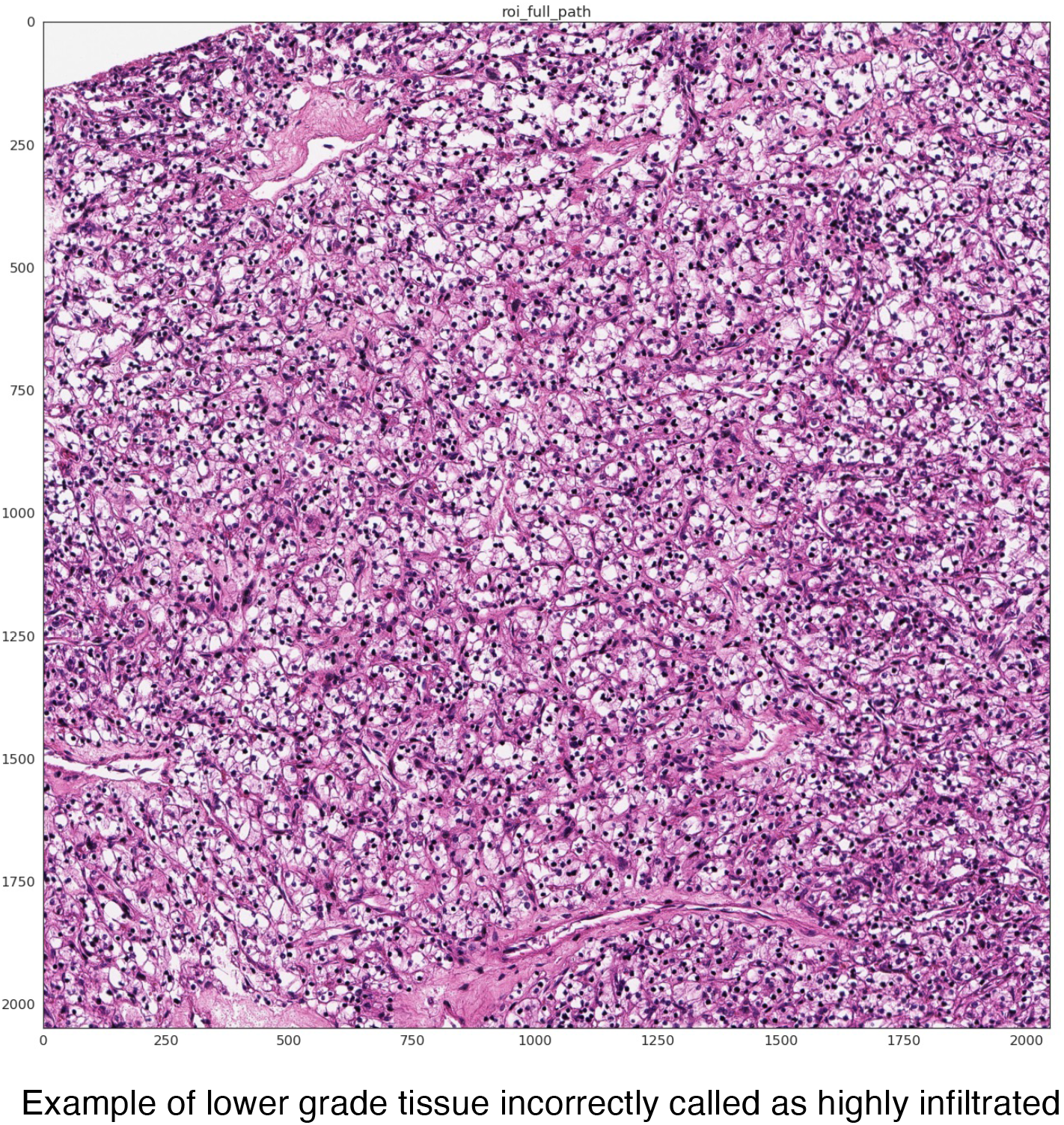
Lower grade tumor tissue is prone to extreme false positive rates, wherein tumor nuclei are classified as TILs incorrectly.

**Extended Data Fig. 6:**
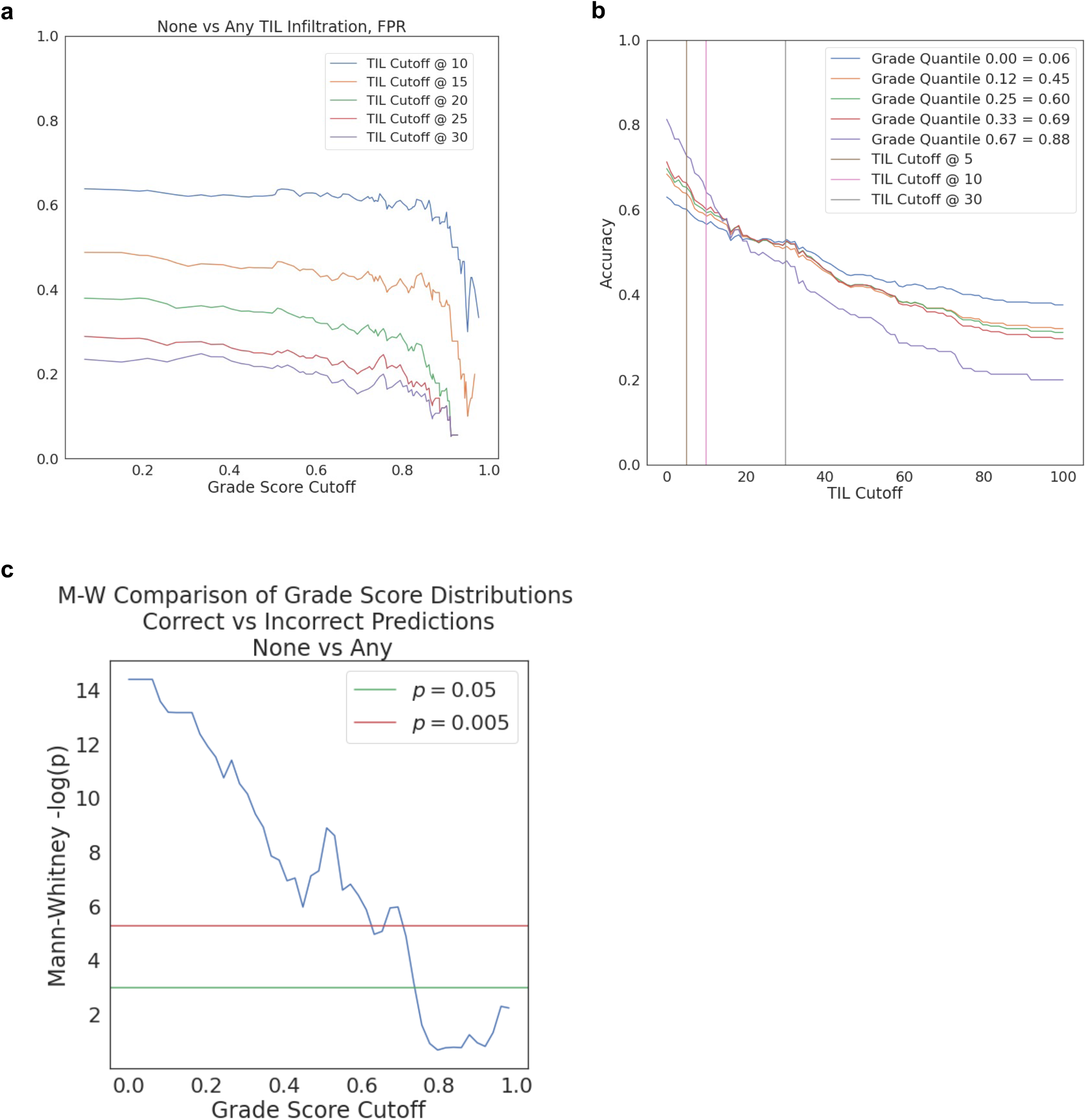
Accuracy in predicting TIL infiltration presence relies on both grade score and infiltration cutoff choice. A/B. False positive rate (FPR) and accuracy, respectively, versus minimum tissue segment grade score required for TIL evaluation at different cutoffs for calling “infiltrated”. C. Mann-Whitney U test statistic p-values from comparing grade score distributions of correctly versus incorrectly classified tiles (“none” vs “any” TIL presence) at different minimum grade score cutoffs.

**Extended Data Fig. 7:**
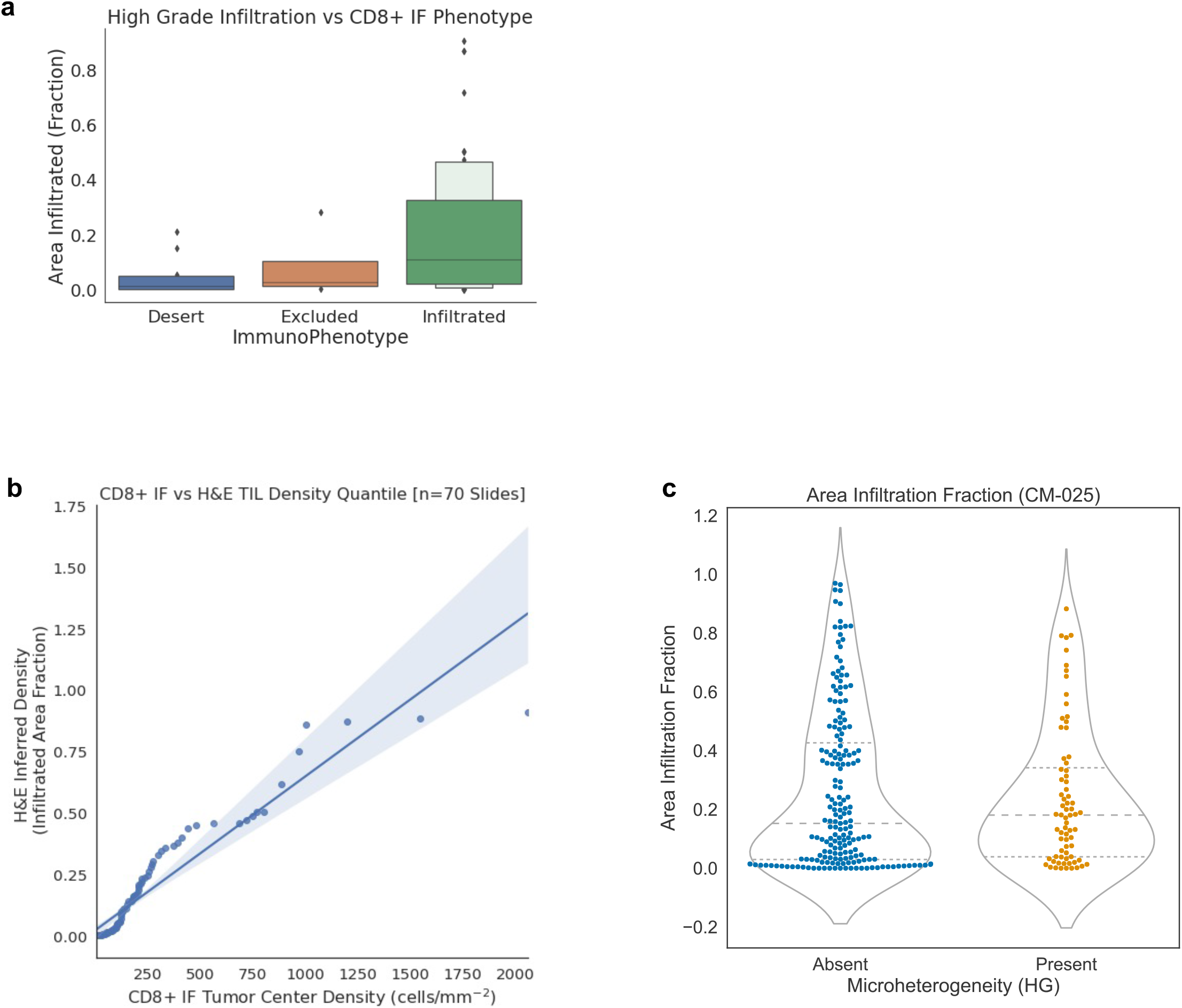
Comparing H&E derived TIL phenotypes to CD8+ data from the same tumor. A. Inferred tumor infiltrating lymphocyte density in high grade foci is consistent with CD8+ immunofluorescence data collected for a subset in the same cohort (Braun et al., 2020). B. QQ-plot comparison of CD8+ IF tumor center cell density versus H&E-inferred TIL infiltrated area fraction. C. Area infiltration fraction in CM-025 versus microheterogeneity status (within edges containing a high grade node [score >= 0.8]). Area infiltration fraction: proportion of tiles above the “infiltrated” cutoff (14 TIL/tile).

**Extended Data Fig. 8:**
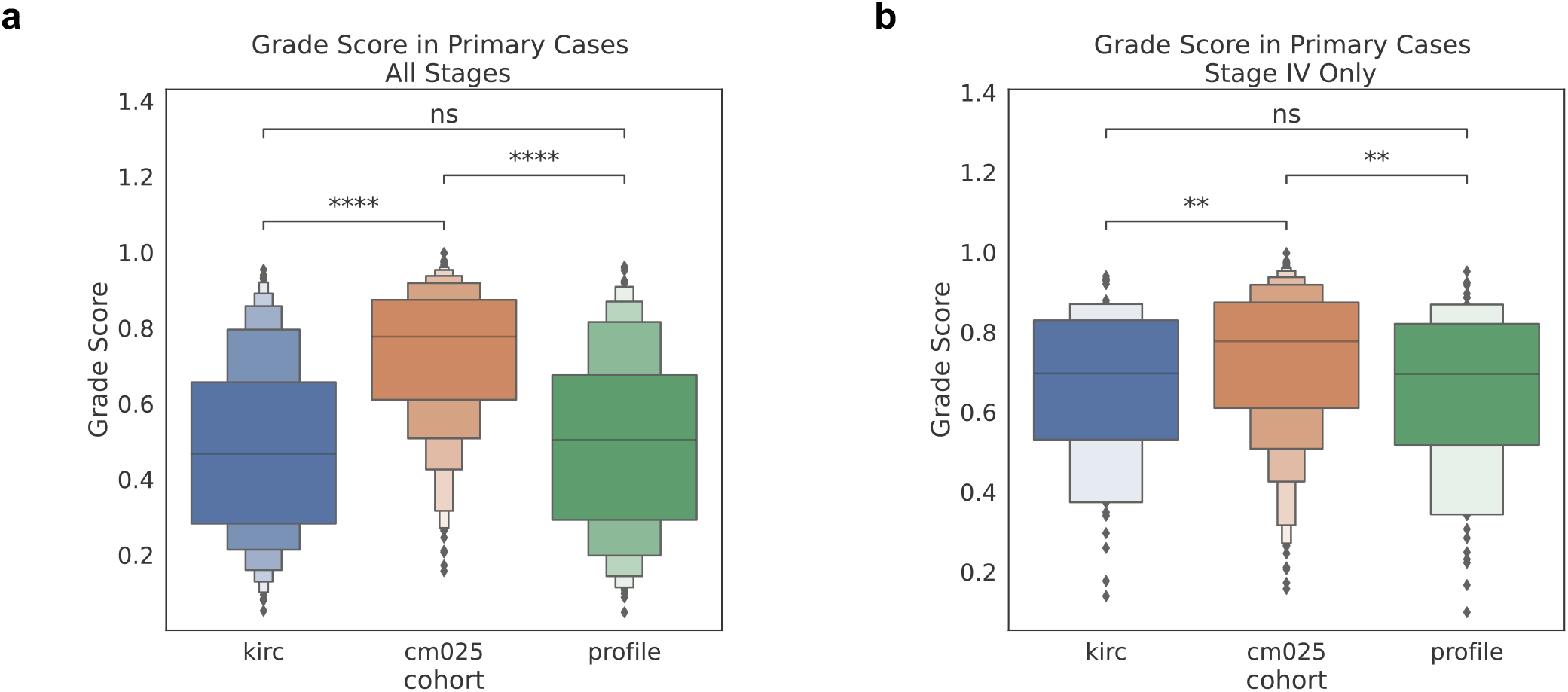
Cohort level distributional differences in grade score. **A.** Grade score distributions for each cohort. **B**. Grade score distributions in each cohort, limited to Stage IV cases. p-value annotation legend: ns: p <= 1.00e+00 *: 1.00e-02 < p <= 5.00e-02 **: 1.00e-03 < p <= 1.00e-02 ***: 1.00e-04 < p <= 1.00e-03 ****: p <= 1.00e-04

**Extended Data Figure 9:**
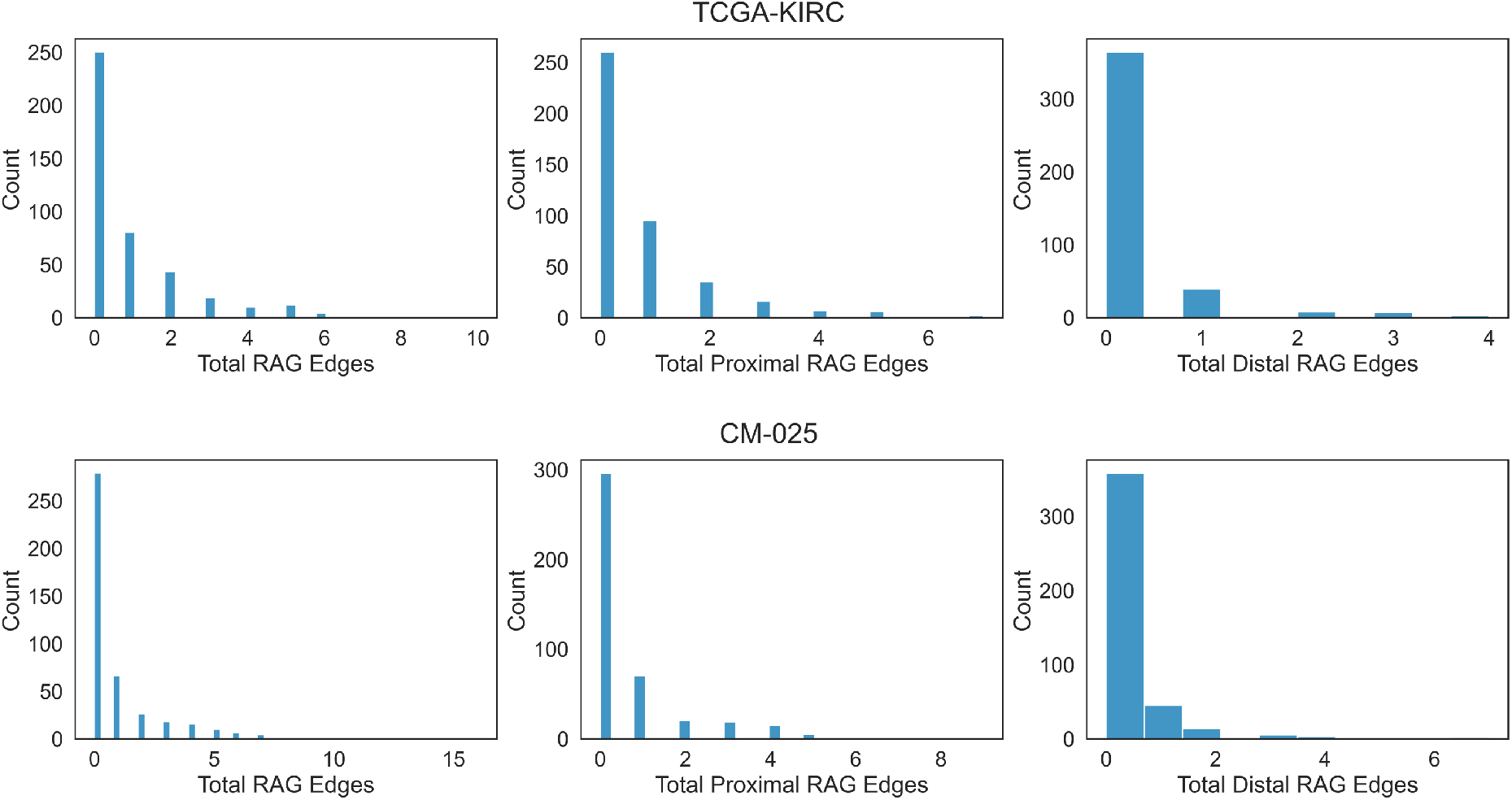
Distribution of RAG edges across TCGA-KIRC and CM-025. Histograms of RAG edge counts, split by type.

**Extended Data Figure 10:**
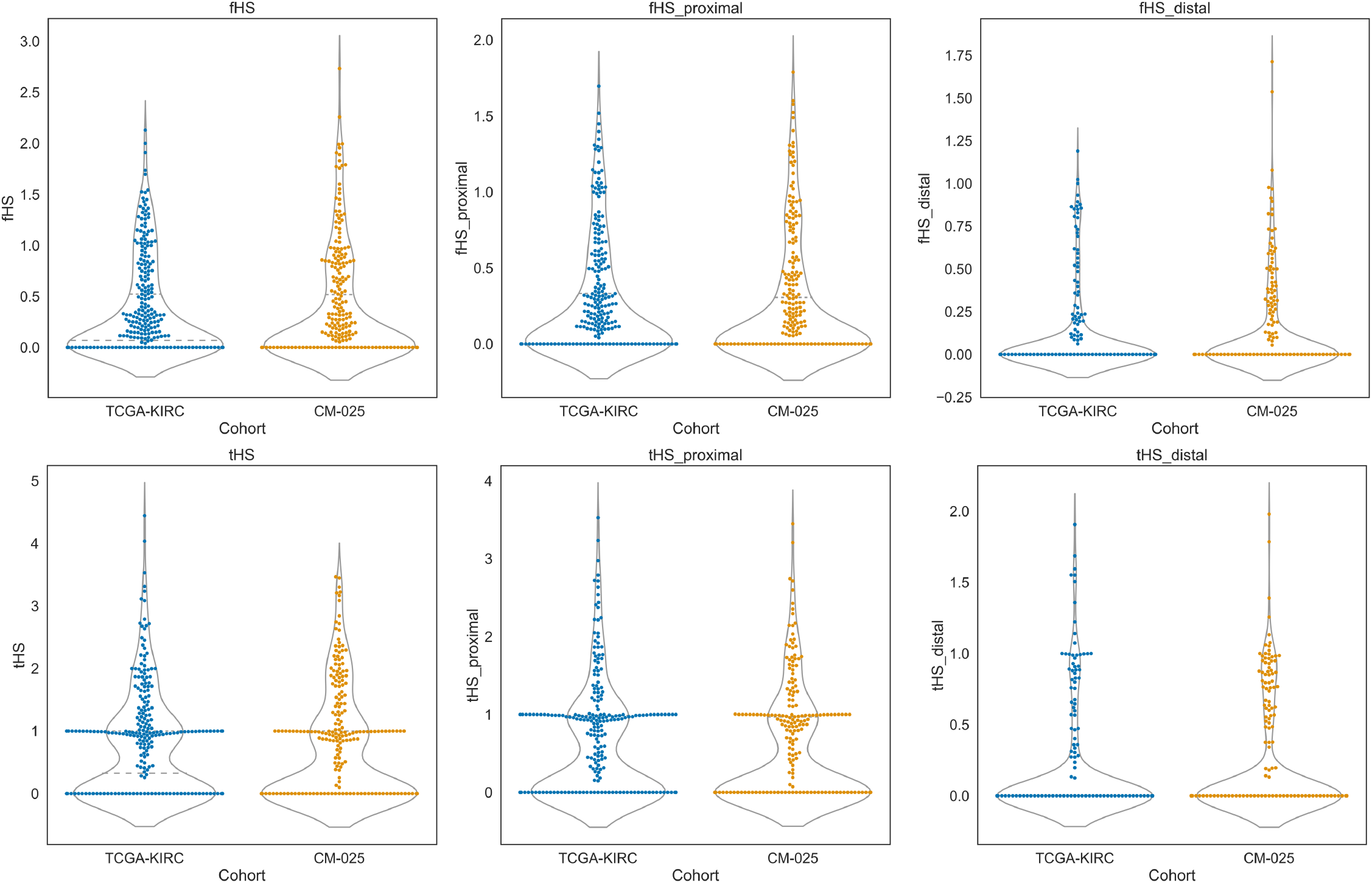
Distribution of continuous heterogeneity scores across TCGA-KIRC and CM-025. Violin-Swarm plots for f-weighted heterogeneity scores (f-HS). Left column: combined (summed) proximal and distal counts/scores. Middle: proximal context (“_proximal”). Right: Distal context (“_distal”).

**Extended Data Figure 11:**
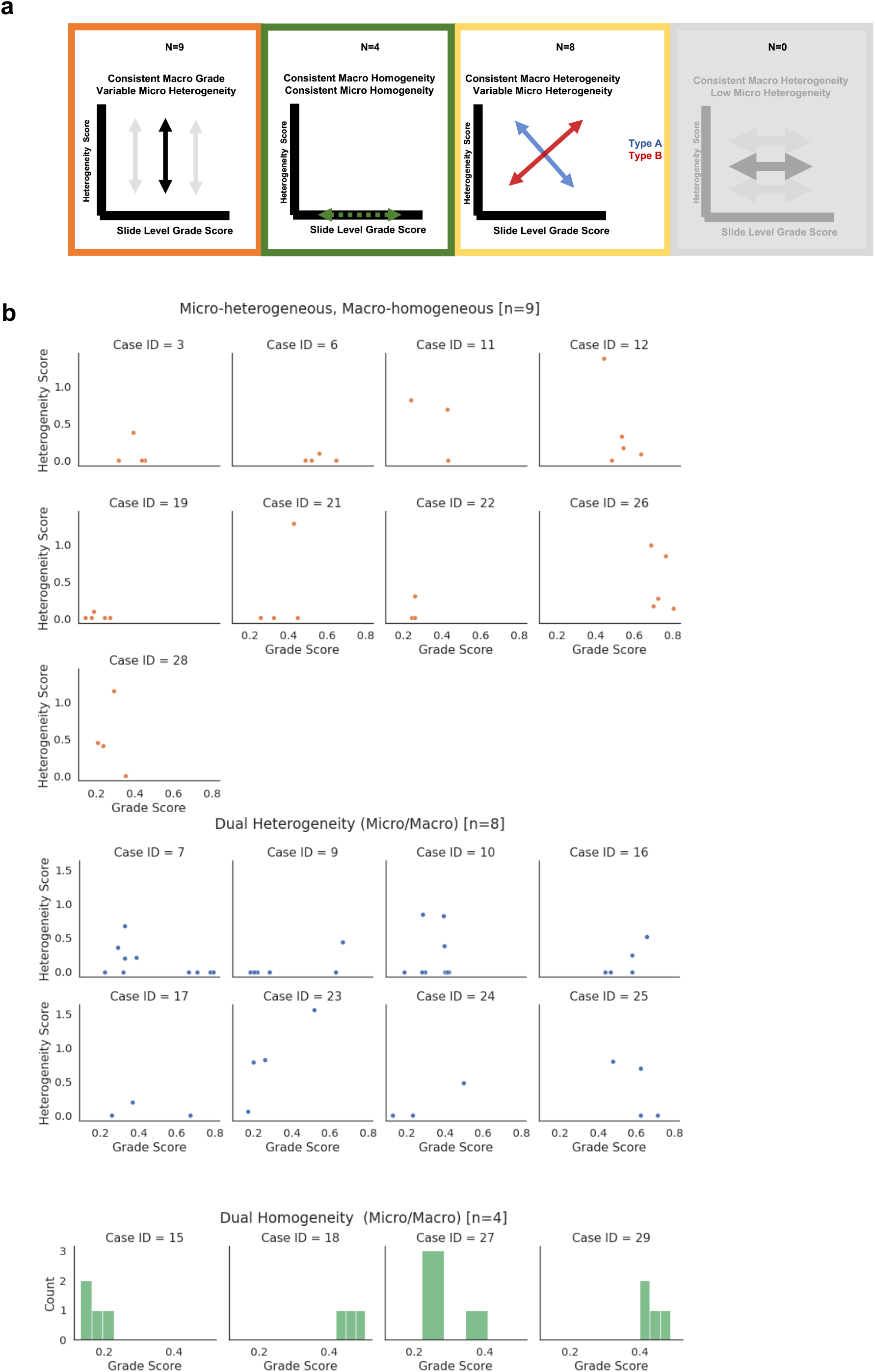
Multiregional microheterogeneity dataset description. A. schematic of proposed covariation patterns of microheterogeneity score and grade score within the samples collected from a single patient tumor. B. Actual data analyzed.

**Extended Data Figure 12:**
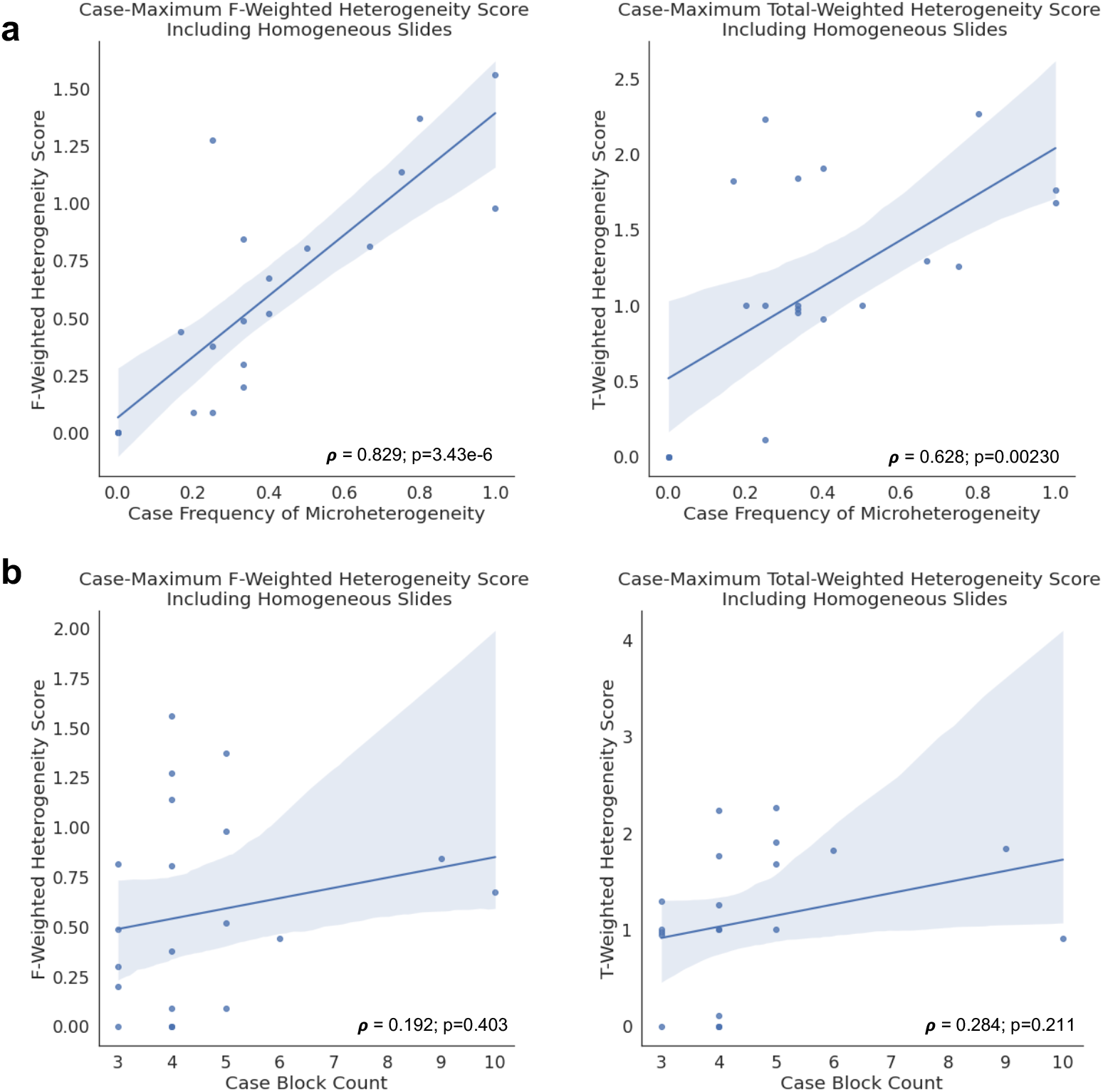
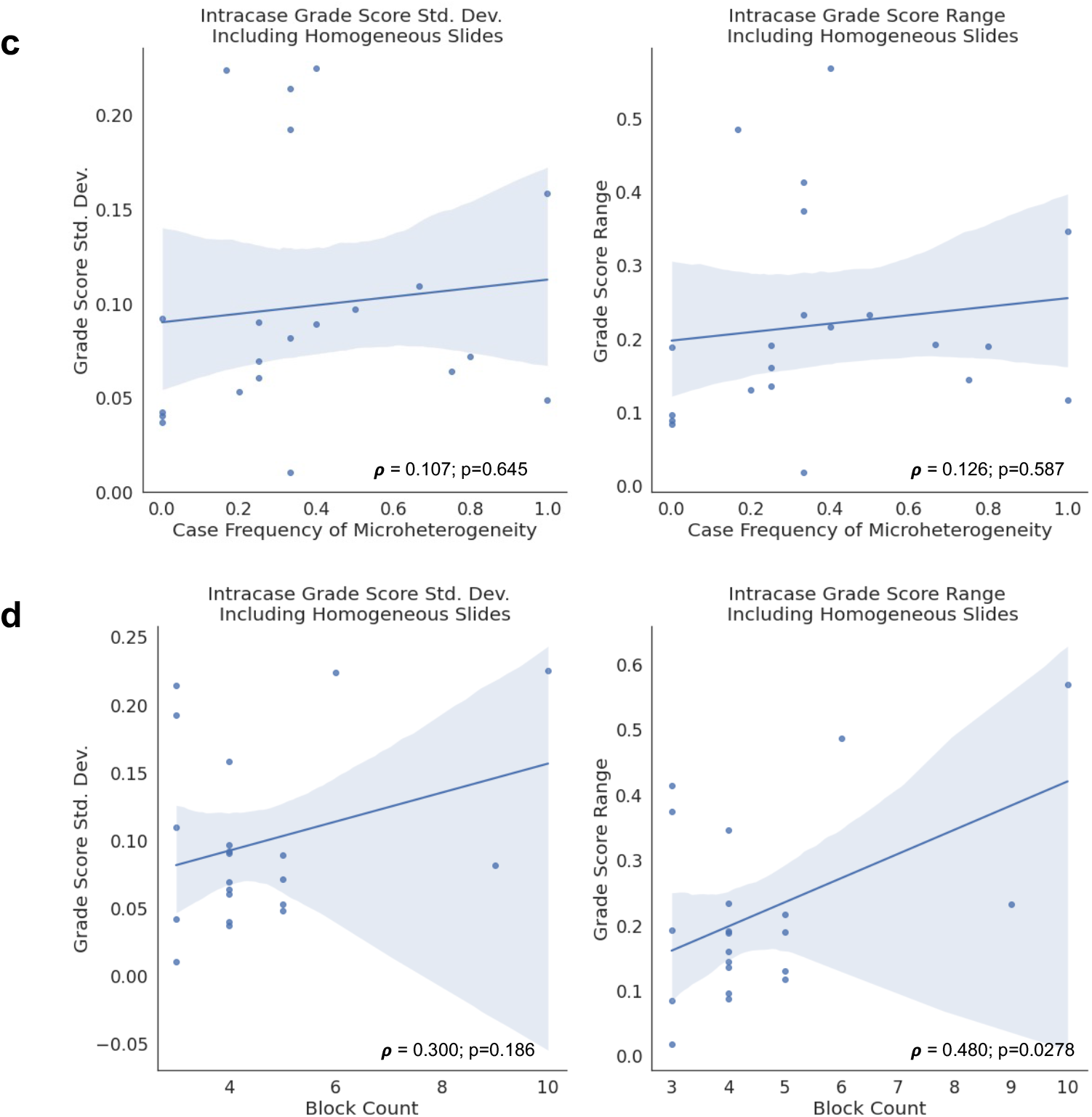
Aggregate continuous heterogeneity scores. A. Case-wise frequency of microheterogeneity versus the maximum observed f-weighted or total-weighted heterogeneity score. B. Case-wise block count versus the maximum observed f-weighted or total-weighted heterogeneity score. Statistics aggregated within a given patient’s set of scanned tissue blocks (1 slide per block). Pearson’s Rho p-values calculated via exact distribution. C. Case-wise frequency of microheterogeneity versus the intracase grade score standard deviation or range. D. Case-wise block count versus intracase grade score standard deviation or range. Statistics aggregated within a given patient’s set of scanned tissue blocks (1 slide per block). Pearson’s Rho p-values calculated via exact distribution.

**Extended Data Figure 13:**
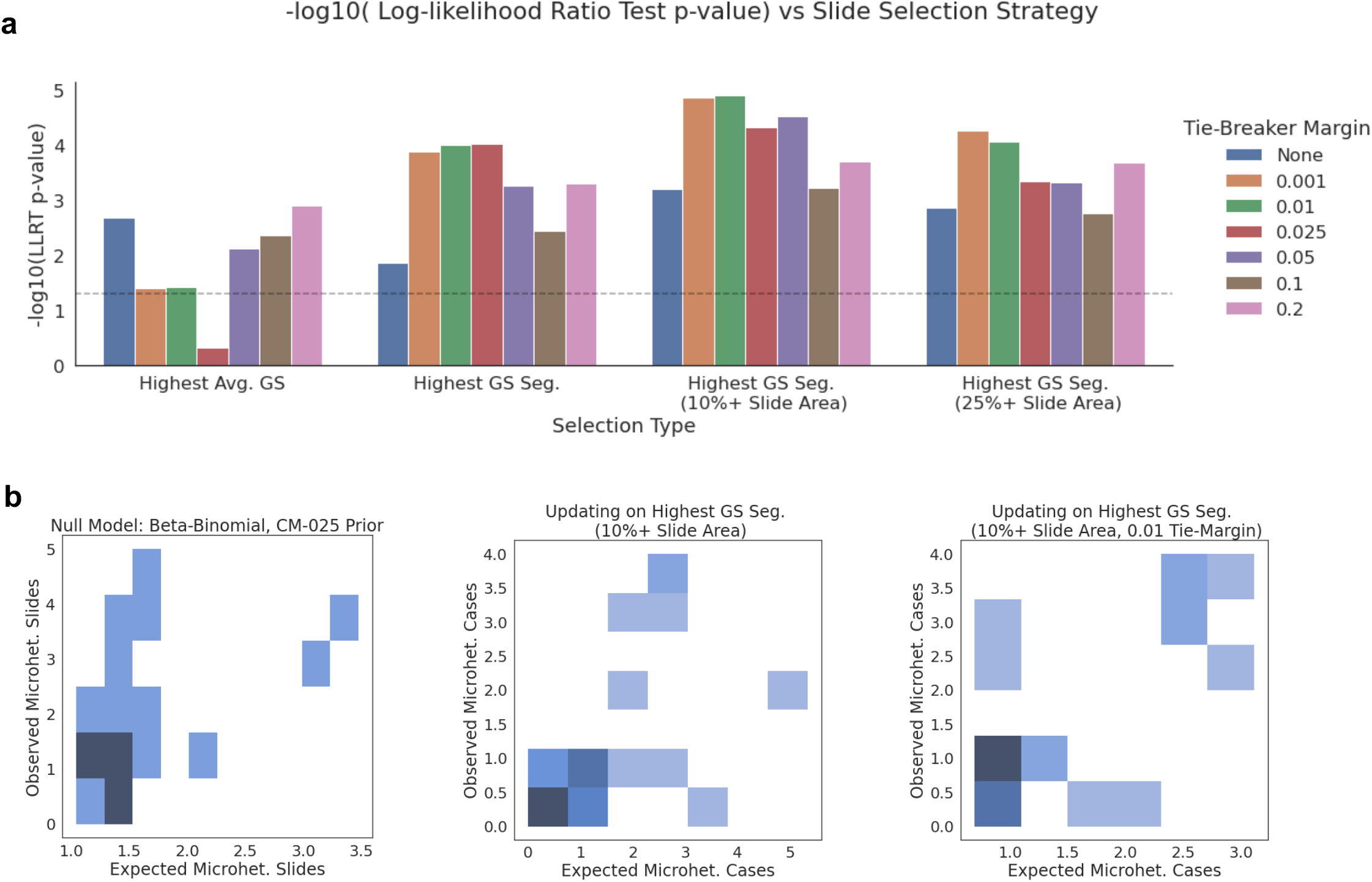
Modeling microheterogeneity occurrence. A. Negative log10 p-values for a likelihood ratio test comparing a null model (beta binomial) to a model that updates its parameters upon observing one “reference” slide from a patient’s collection of samples. X-axis: different selection strategies. Tie-Breaker Margin: margin to allow for considering two or more samples as equally weighted references. GS: Grade Score. B. visualization of expected versus observed data for predicting occurrence of microheterogeneity based on null model or alternative reference model strategy.

**Extended Data Figure 14:**
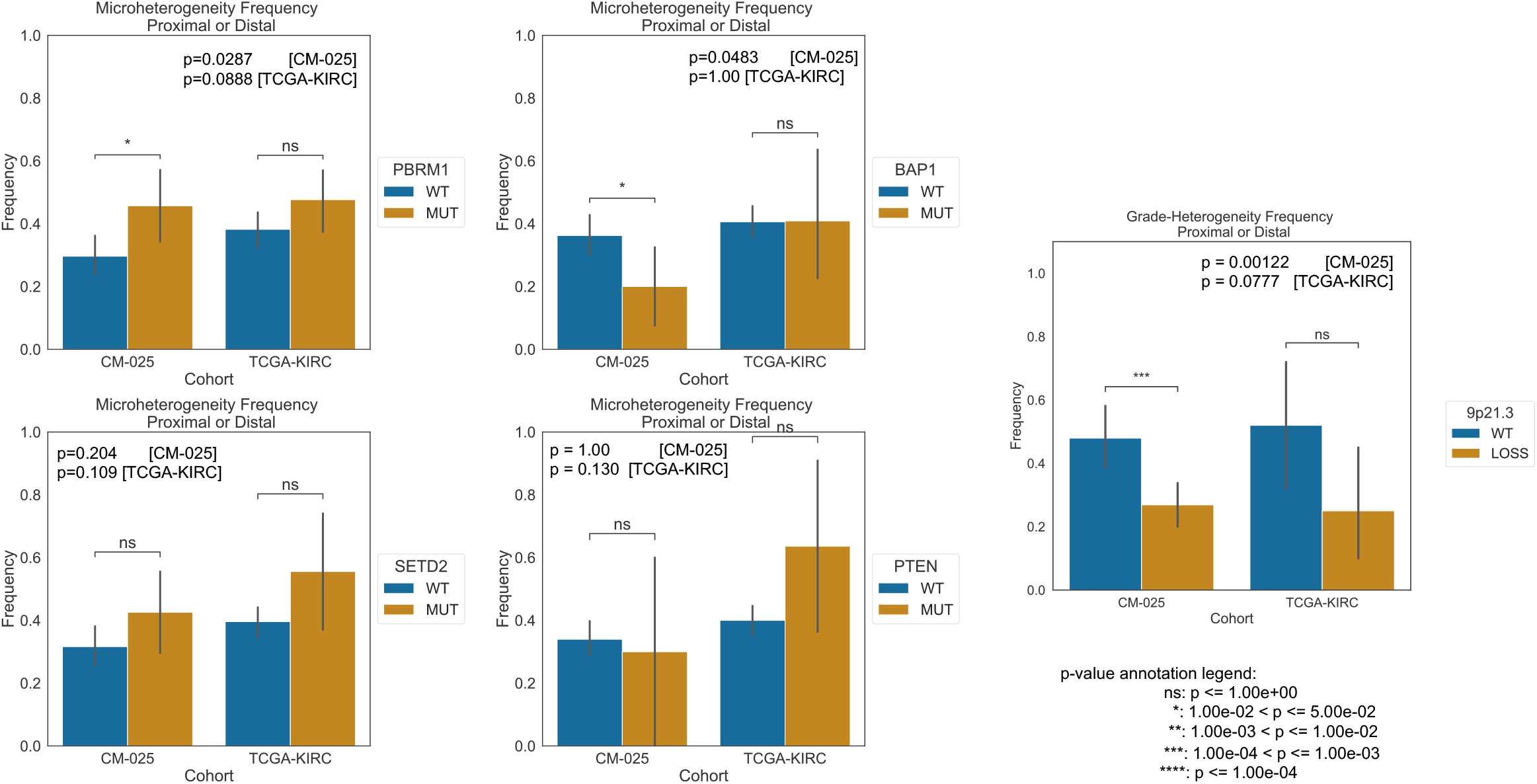
Frequency of microheterogeneity within different loss of function states in TCGA-KIRC and CM-025. Significance calculated with Fisher’s Exact test.

**Extended Data Figure 15.**
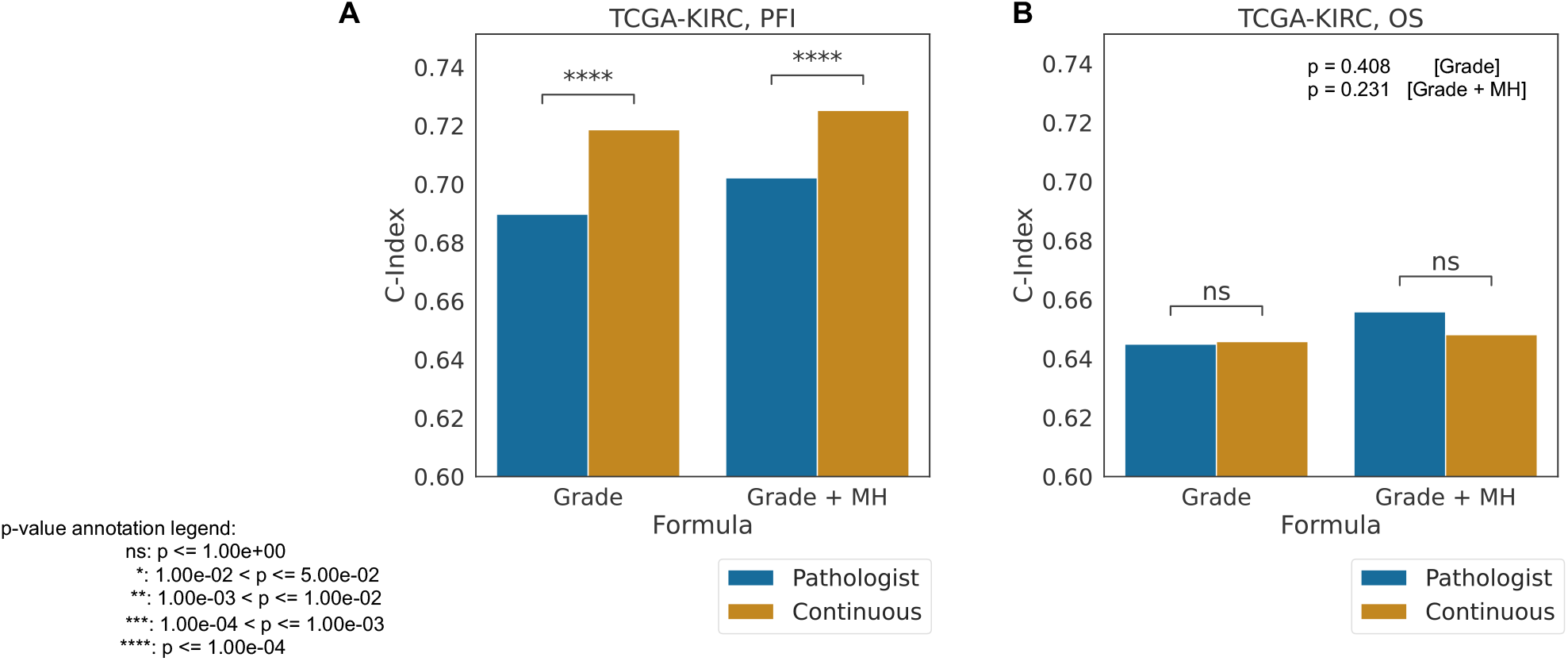
Prognostic correlates of microheterogeneity. Concordance Index (C-Index) for univariate and bivariate models of PFI and OS in TCGA-KIRC. “Grade”: grade type (pathologist vs continuous). “MH” microheterogeneity status (present/absent) as second included covariate. Significance calculated via relative likelihood.

**Extended Data Figure 16:**
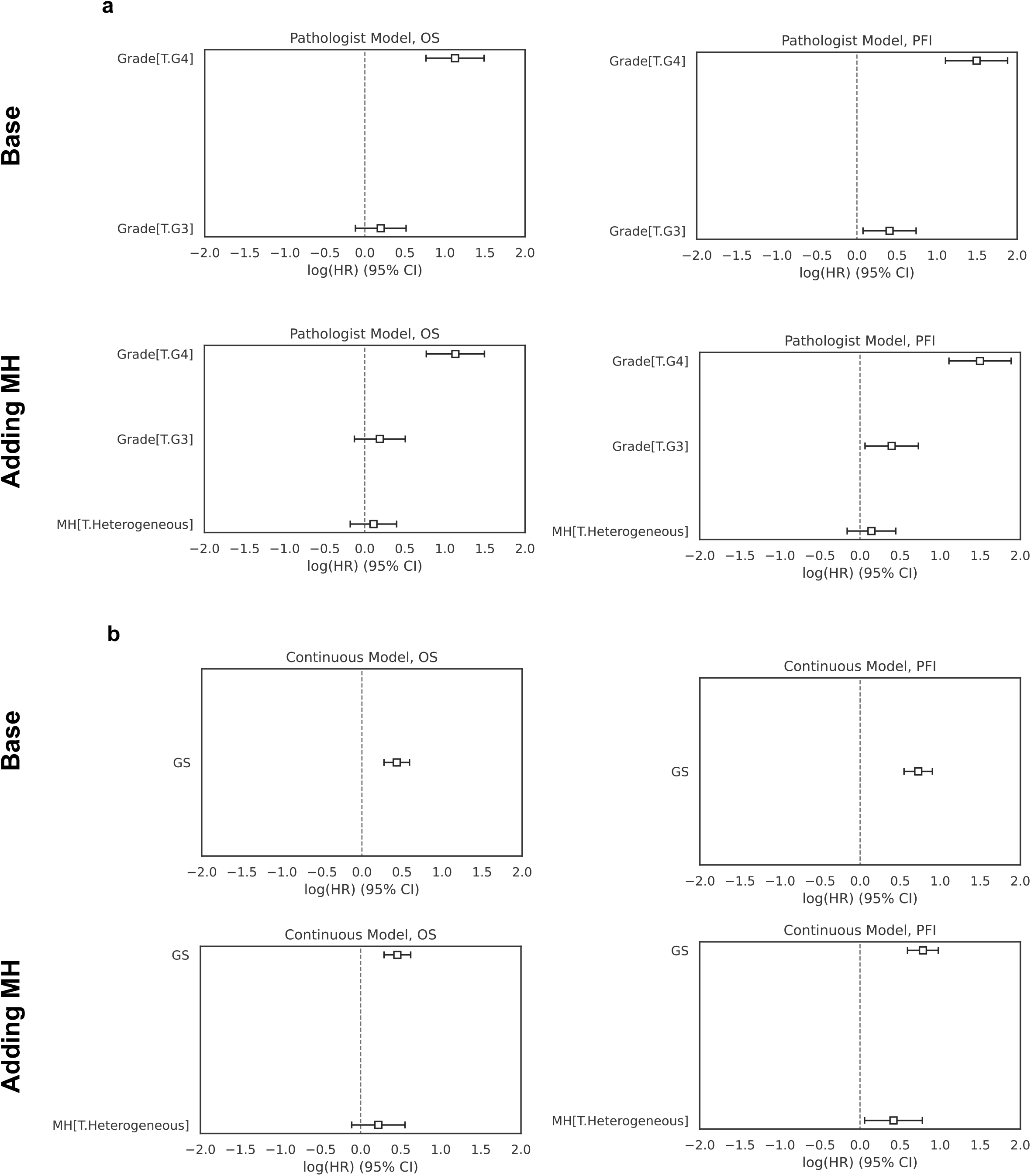
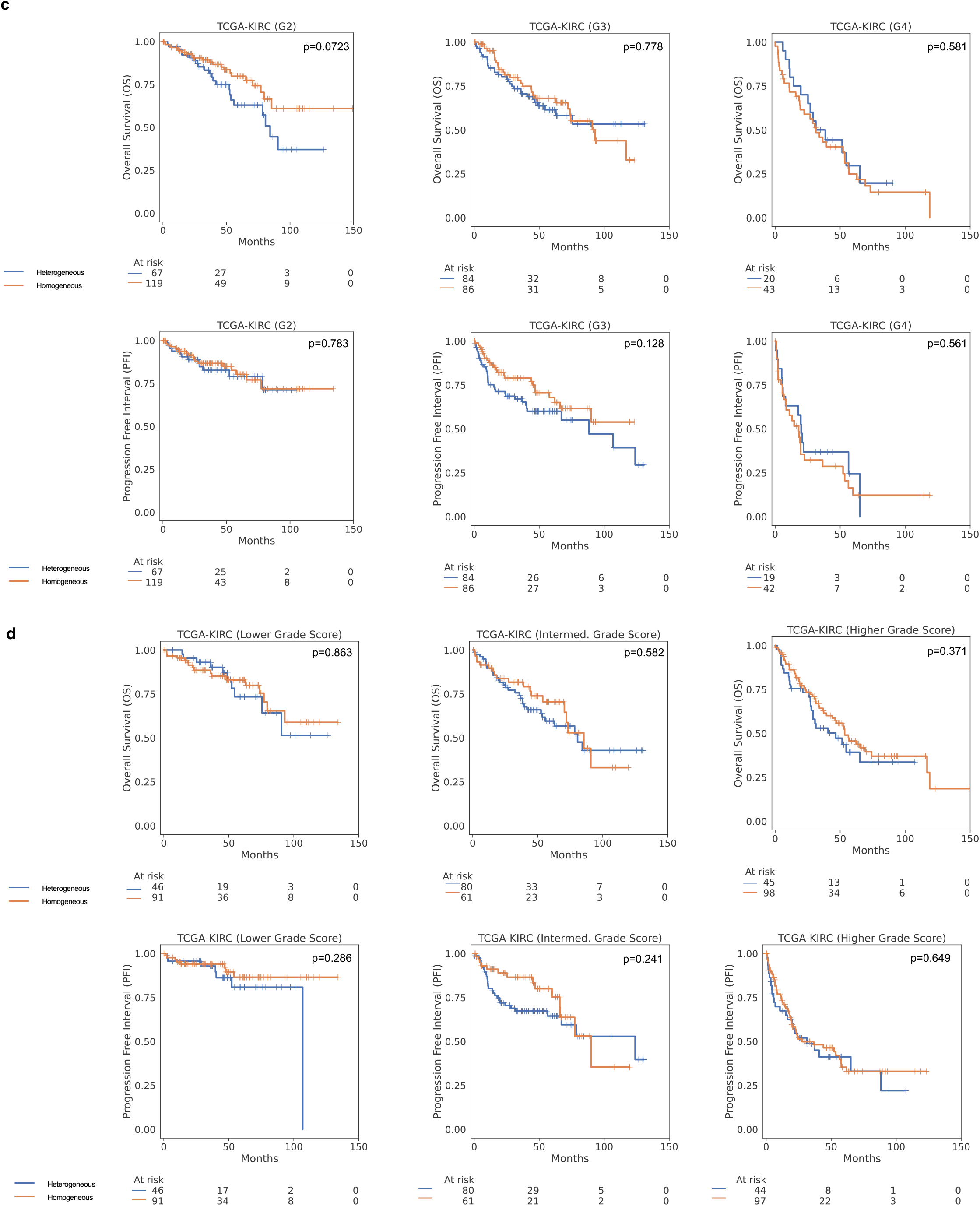
Coefficients for univariate and bivariate Cox proportional hazards models for OS/PFI in TCGA-KIRC. A: pathologist grade. B: continuous grade. “Base”: single covariate type(s). “MH”: microheterogeneity binary presence. C: pathologist grade. D: continuous grade. P-values calculated via log-rank test.

**Extended Data Fig. 17:**
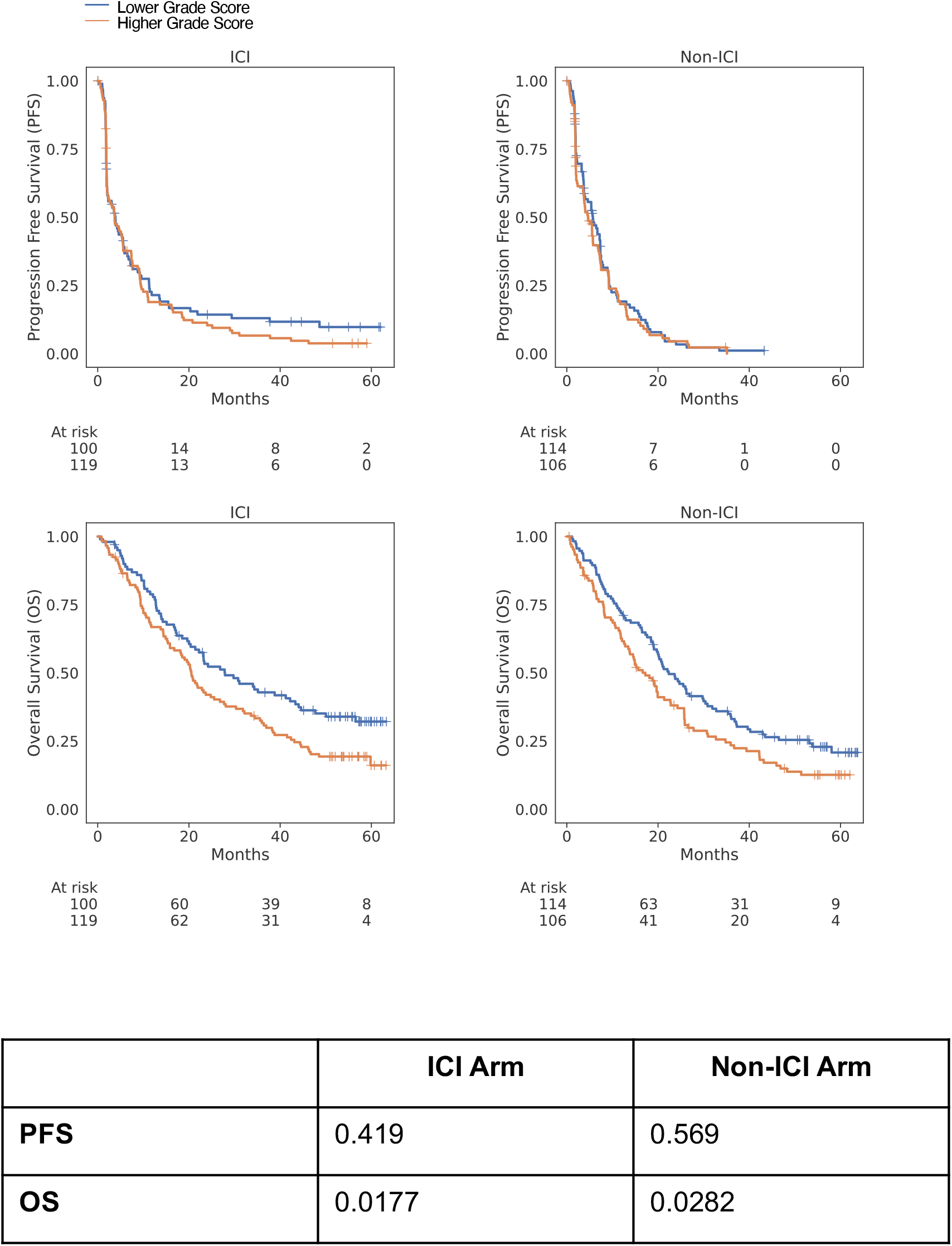
Kaplan-Meier curves for low versus high grade score within each arm of the CM-025 trial. Top row: progression free survival (PFS). Bottom row: overall survival (OS). Stratification based on the median inferred grade score in the CM-025 cohort. Table: log-rank test p-values for shown curves.

**Extended Data Fig. 18:**
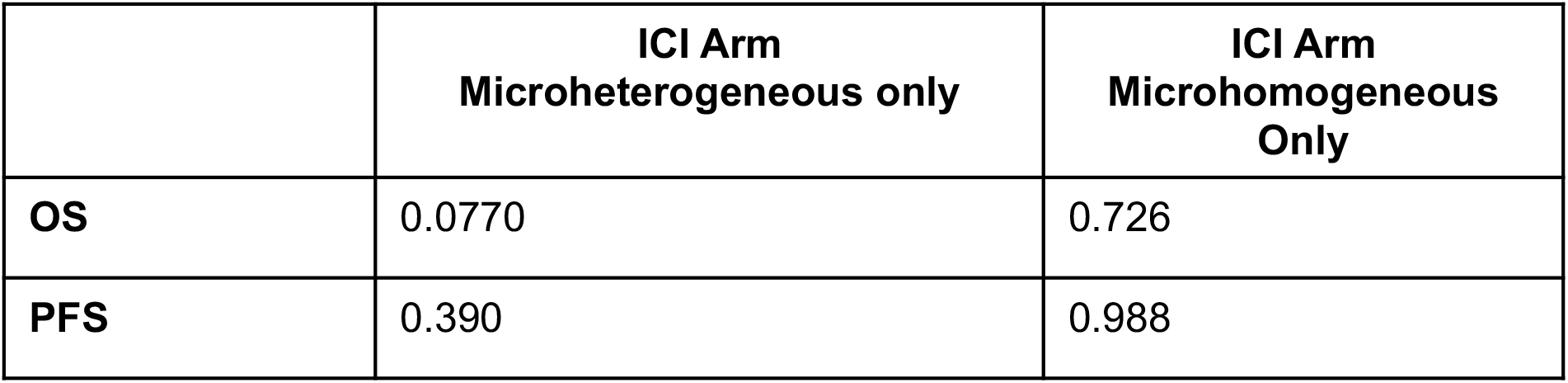
Likelihood ratio test evaluation of univariate Cox proportional hazards models using continuous grade score in the ICI arm of CM-025. Columns: subset of data; rows: survival endpoint modelled.

**Extended Data Fig. 19:**
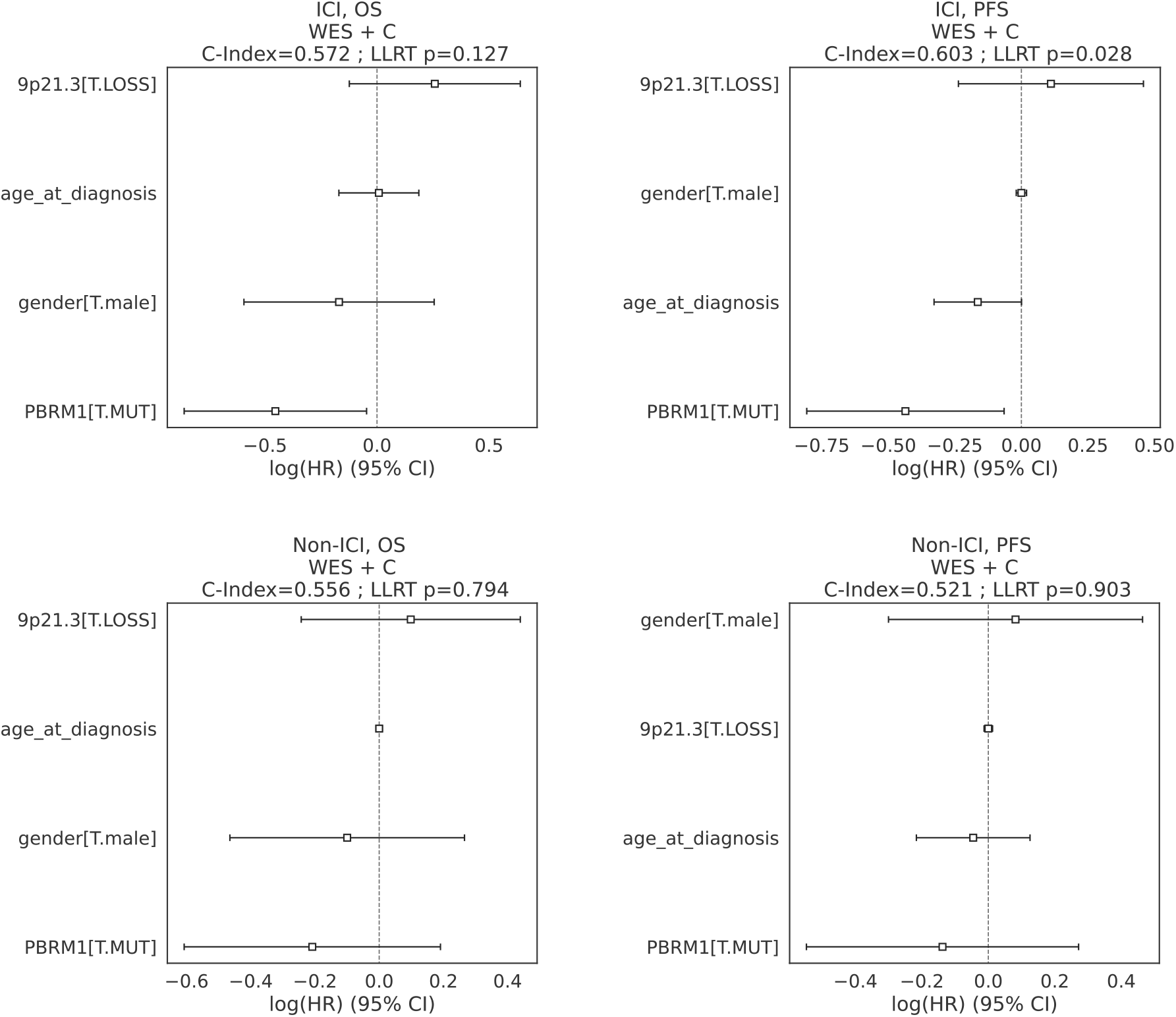
Cox model coefficients for models in the CM-025 cohort, limited to genomic and clinical features (“WES + C”). LLRT: loglikelihood ratio test. C-Index: concordance index. WES: whole-exome sequencing.

**Extended Data Fig. 20:**
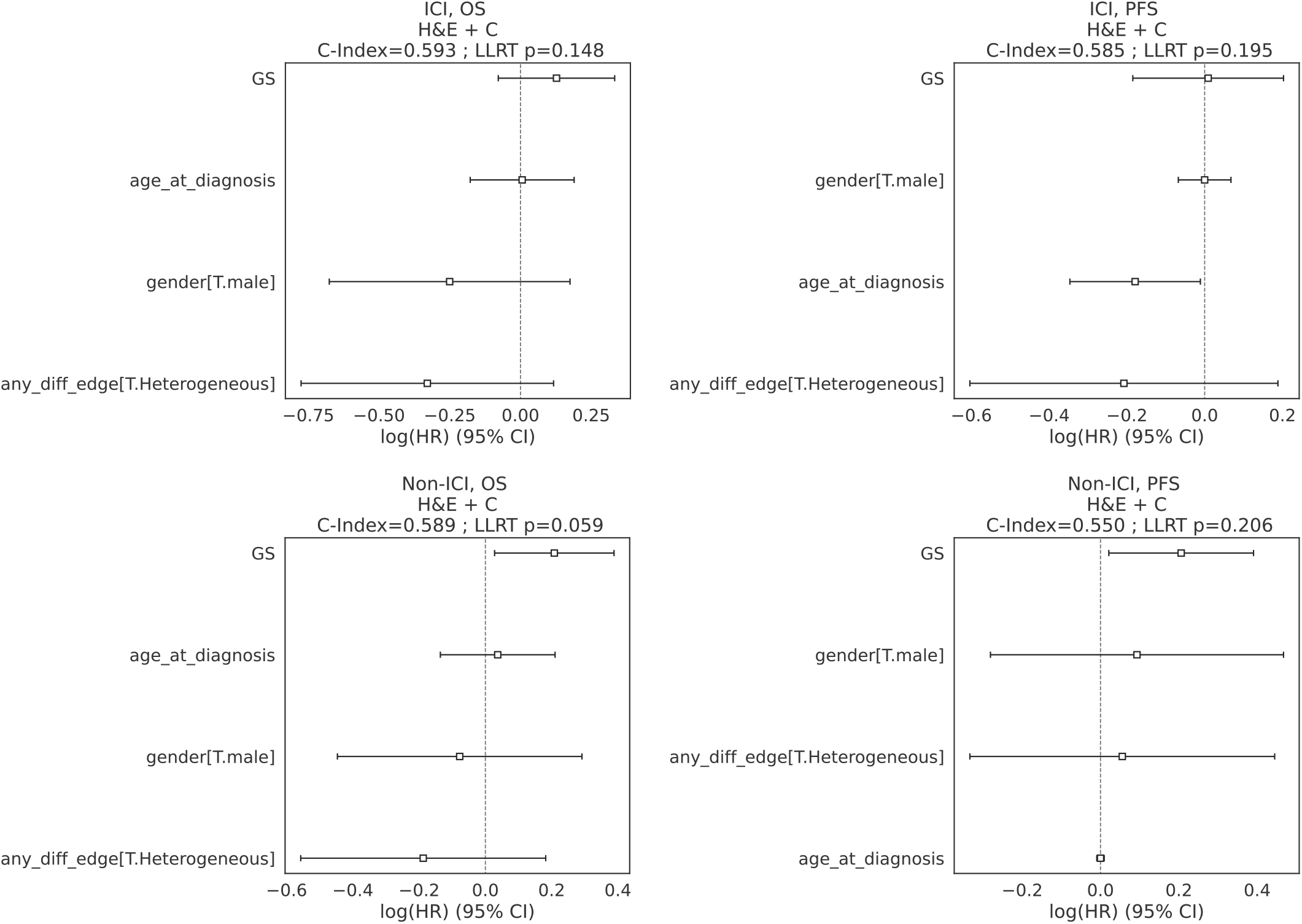
Cox model coefficients for models in the CM-025 cohort, limited to H&E/computer vision and clinical features (“H&E + C”). LLRT: loglikelihood ratio test. C-Index: concordance index. ‘any_diff_edge’: microheterogeneity categorical variable. GS: continuous grade score.

**Extended Data Fig. 21:**
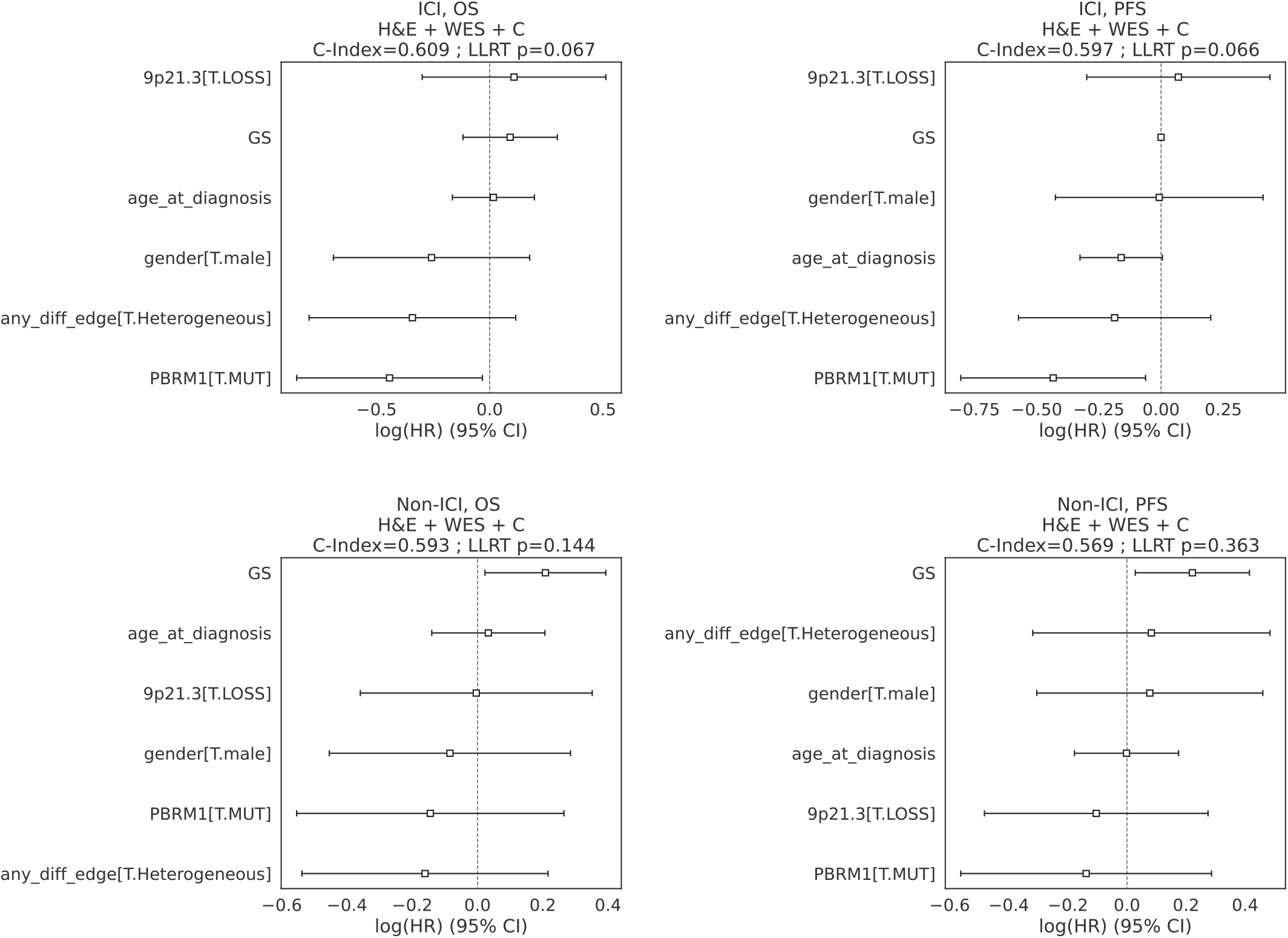
Cox model coefficients for models in the CM-025 cohort, limited to genomic, H&E/computer vision and clinical features (“H&E + WES + C”). LLRT: loglikelihood ratio test. C-Index: concordance index. ‘any_diff_edge’: microheterogeneity categorical variable. WES: whole-exome sequencing. GS: continuous grade score.

**Extended Data Fig. 22:**
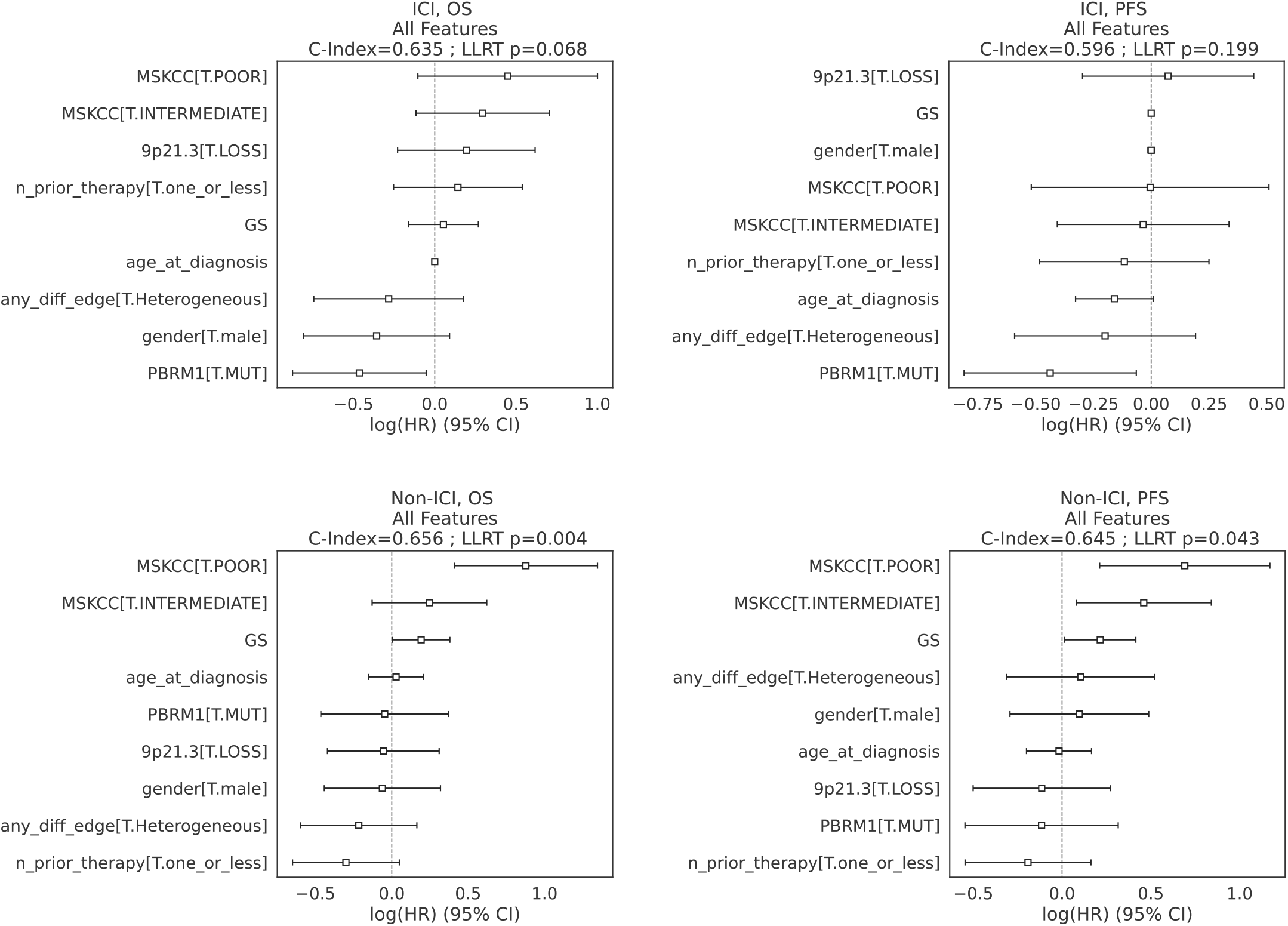
Cox model coefficients for models in the CM-025 cohort, using all available covariate types (genomic, H&E/computer vision, clinical, risk). LLRT: loglikelihood ratio test. C-Index: concordance index. ‘any_diff_edge’: microheterogeneity categorical variable. GS: continuous grade score. MSKCC: MSKCC risk group (categorical).’n_prior_therapy’: number of lines of therapies administered prior to the trial.

**Extended Data Fig. 23:**
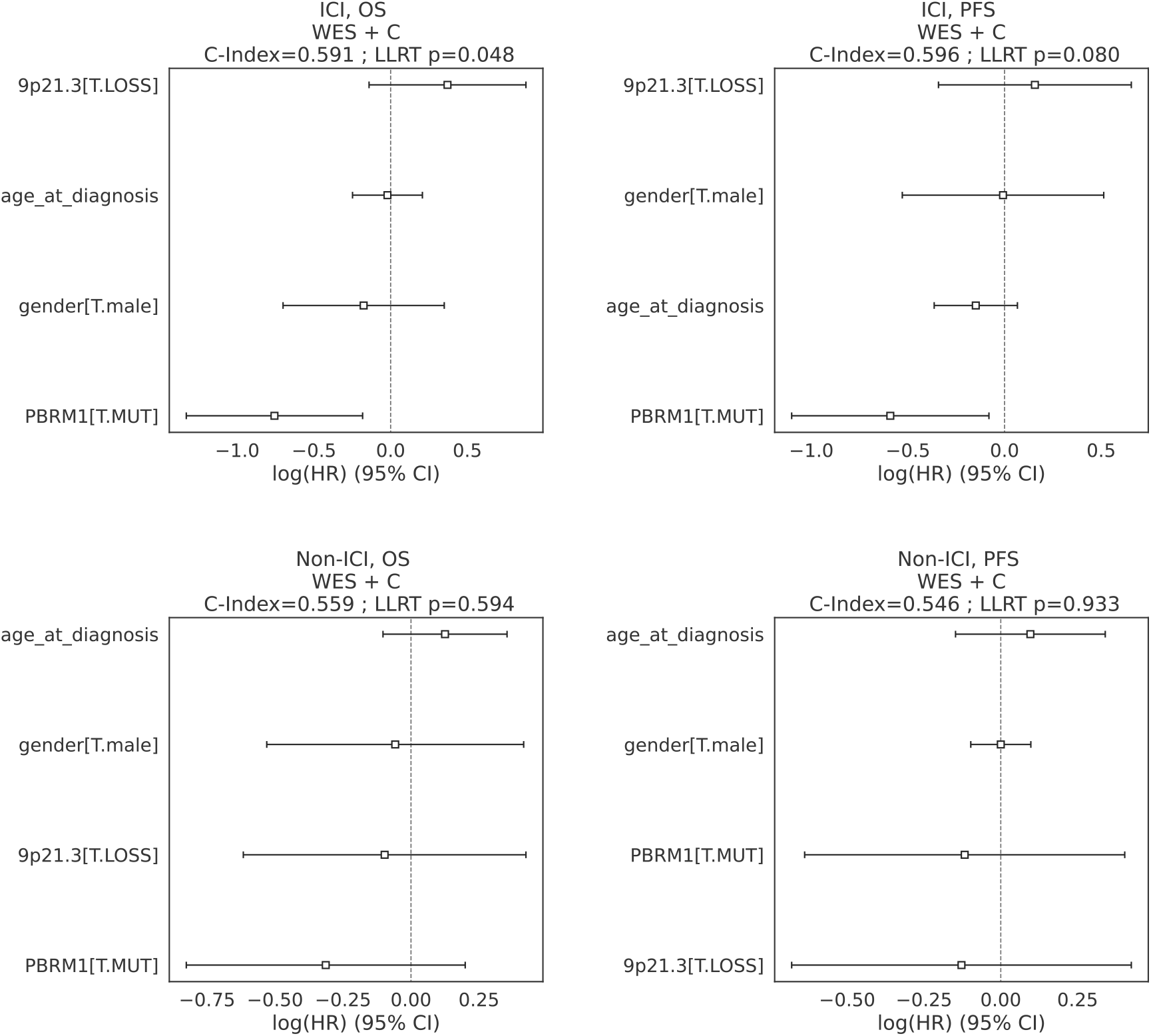
Cox model coefficients for models in the CM-025 cohort, limited to genomic and clinical features, restricted to subset where TIL are evaluable (“WES + C”). LLRT: loglikelihood ratio test. C-Index: concordance index. ‘any_diff_edge’: microheterogeneity categorical variable.

**Extended Data Fig. 24:**
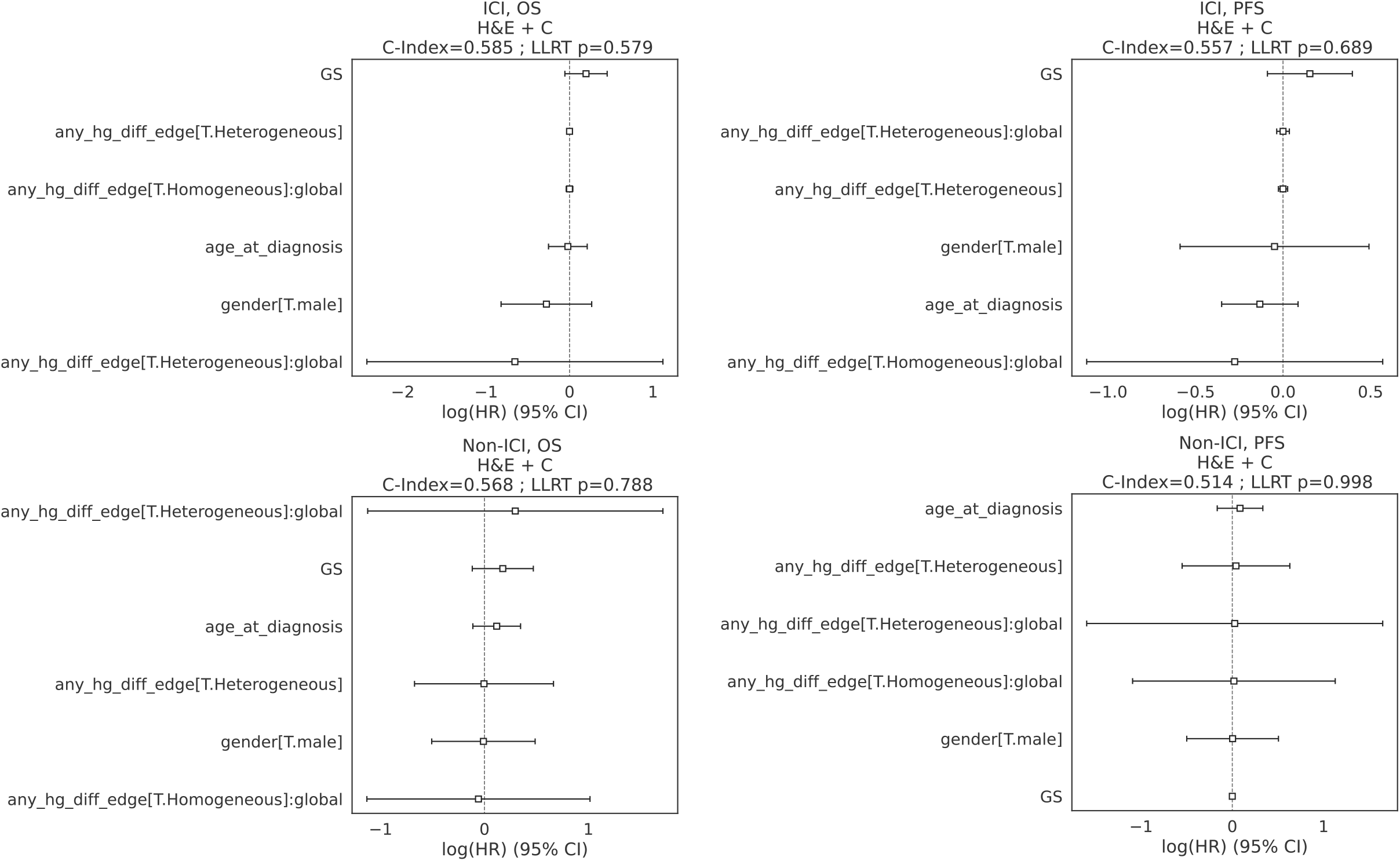
Cox model coefficients for models in the CM-025 cohort, limited to H&E/computer vision [TIL included] and clinical features (“H&E + C”). LLRT: loglikelihood ratio test. C-Index: concordance index. ‘any_hg_diff_edge’: microheterogeneity categorical variable (high-grade node involved in RAG edge required). GS: continuous grade score. Global: area infiltration fraction across evaluated tumor area (fraction tiles above minimum TIL count).

**Extended Data Fig. 25:**
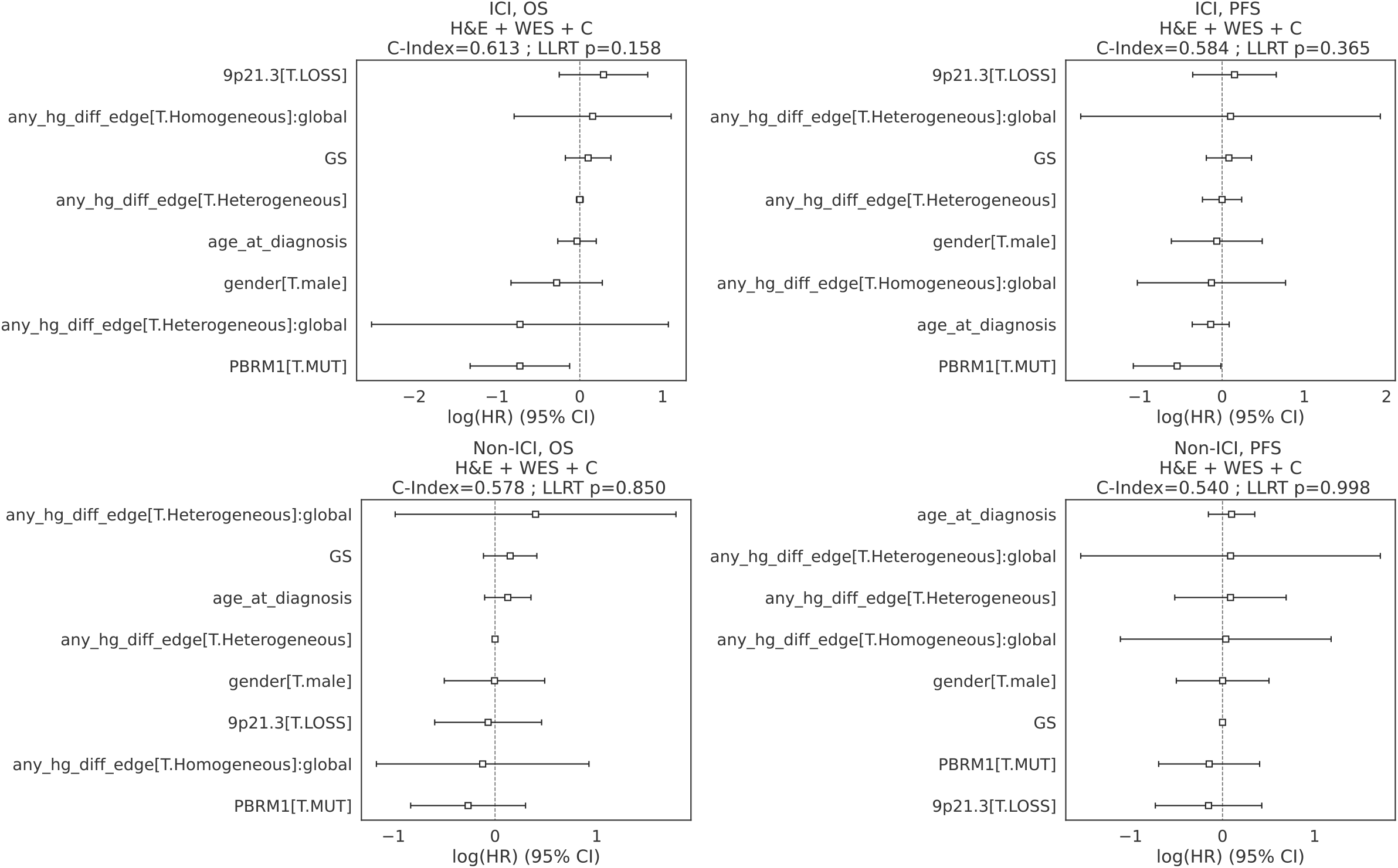
Cox model coefficients for models in the CM-025 cohort, limited to genomic, H&E/computer vision [TIL included] and clinical features (“H&E + WES + C”). LLRT: loglikelihood ratio test. C-Index: concordance index. ‘any_hg_diff_edge’: microheterogeneity categorical variable (high-grade node involved in RAG edge required). GS: continuous grade score. Global: area infiltration fraction across evaluated tumor area (fraction tiles above minimum TIL count).

**Extended Data Fig. 26:**
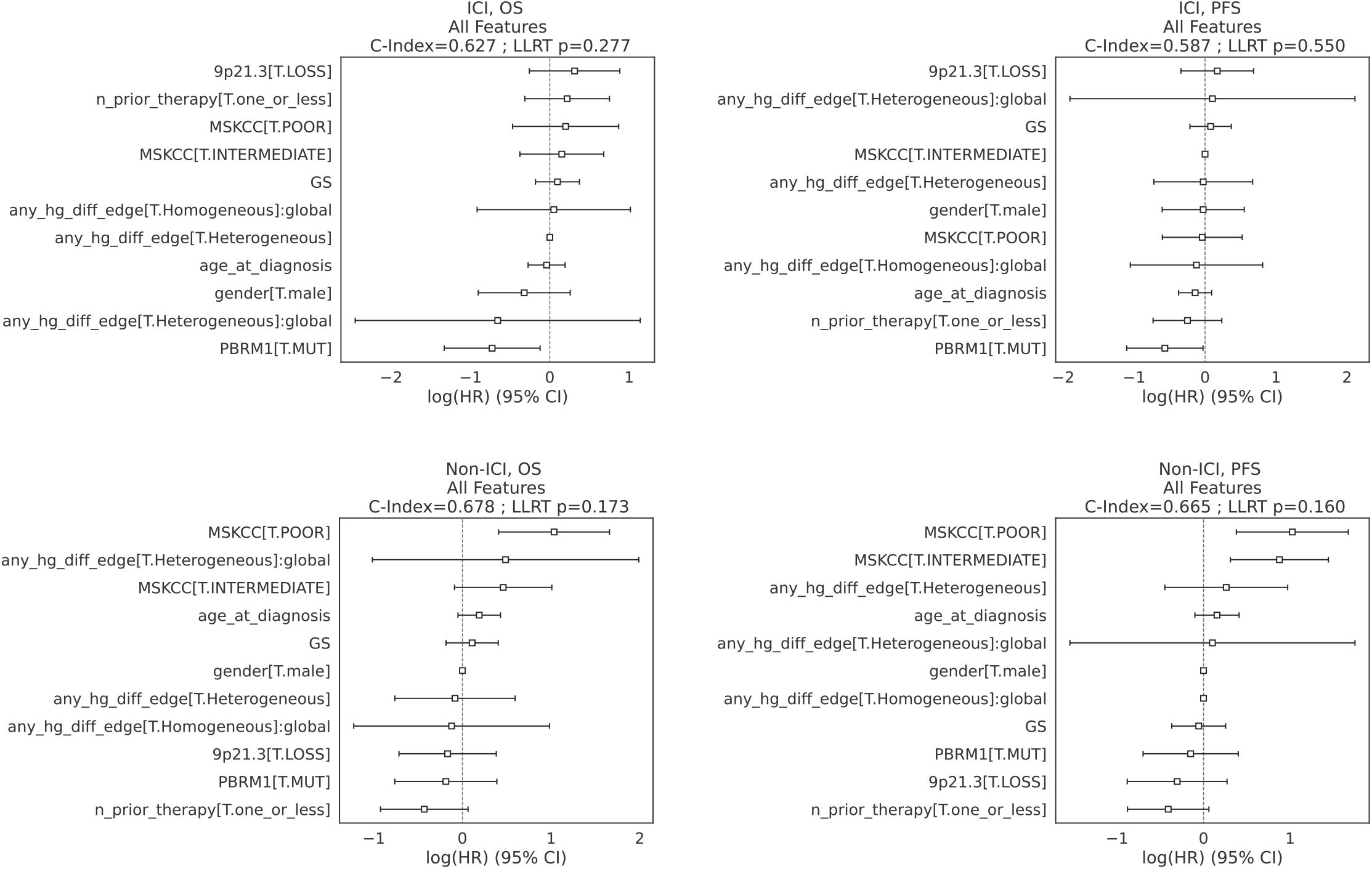
Cox model coefficients for models in the CM-025 cohort, using all available covariate types (genomic, H&E/computer vision [TIL included], clinical, risk). LLRT: loglikelihood ratio test. C-Index: concordance index. ‘any_hg_diff_edge’: microheterogeneity categorical variable (high-grade node involved in RAG edge required). GS: continuous grade score. Global: area infiltration fraction across evaluated tumor area (fraction tiles above minimum TIL count). MSKCC: MSKCC risk group (categorical). ‘n_prior_therapy’: number of lines of therapies administered prior to the trial.

**Extended Data Figure 27:**
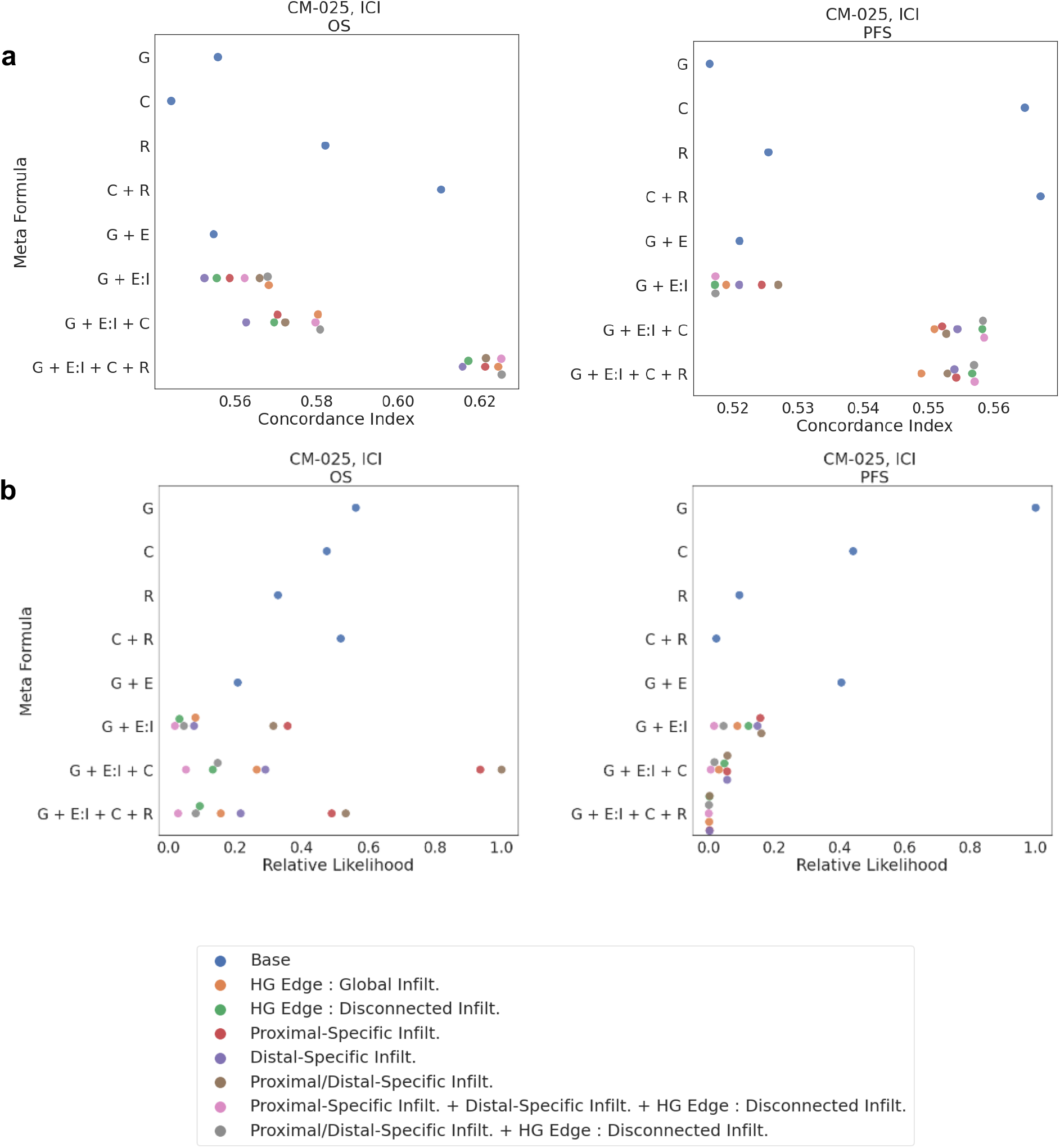
Comparison of different immune-context specifications when fitting Cox proportional hazards models for overall survival in CM-025, ICI arm. A. Concordance Index. B. Relative Likelihood. Colors indicate the form of infiltration covariate used. HG Edge: edge involving a high-grade node (average score above 0.8). *Global*: Using all evaluable tumor area for infiltration fraction description. *Disconnected*: Using nodes that are disconnected from RAG. *Proximal* or *Distal*: infiltration specific to a proximal or distal edge, respectively. G: <u>G</u>rade Score E: Heterogeneity/RAG <u>E</u>dge Variable I: Infiltration Variable C: <u>C</u>linical Base Info (Sex, Age) R: Clinical <u>R</u>isk/Performance Info (MSKCC Risk group, Num. prior lines Tx before trial) (E:I is shorthand for [E + E*I], where E:I is an interaction term between variables E and I)

**Extended Data Fig. 28:**
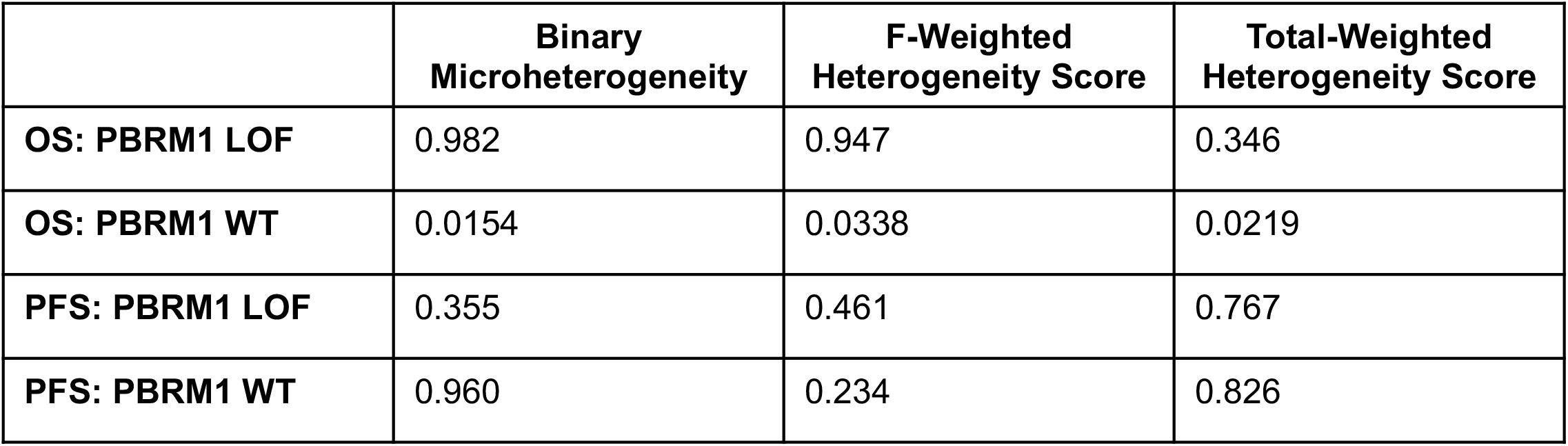
Comparison of Cox model LLRT p-values under different PBRM1 states in the ICI arm of CM-025. LOF: loss of function (truncating mutation present).

**Extended Data Figure 29:**
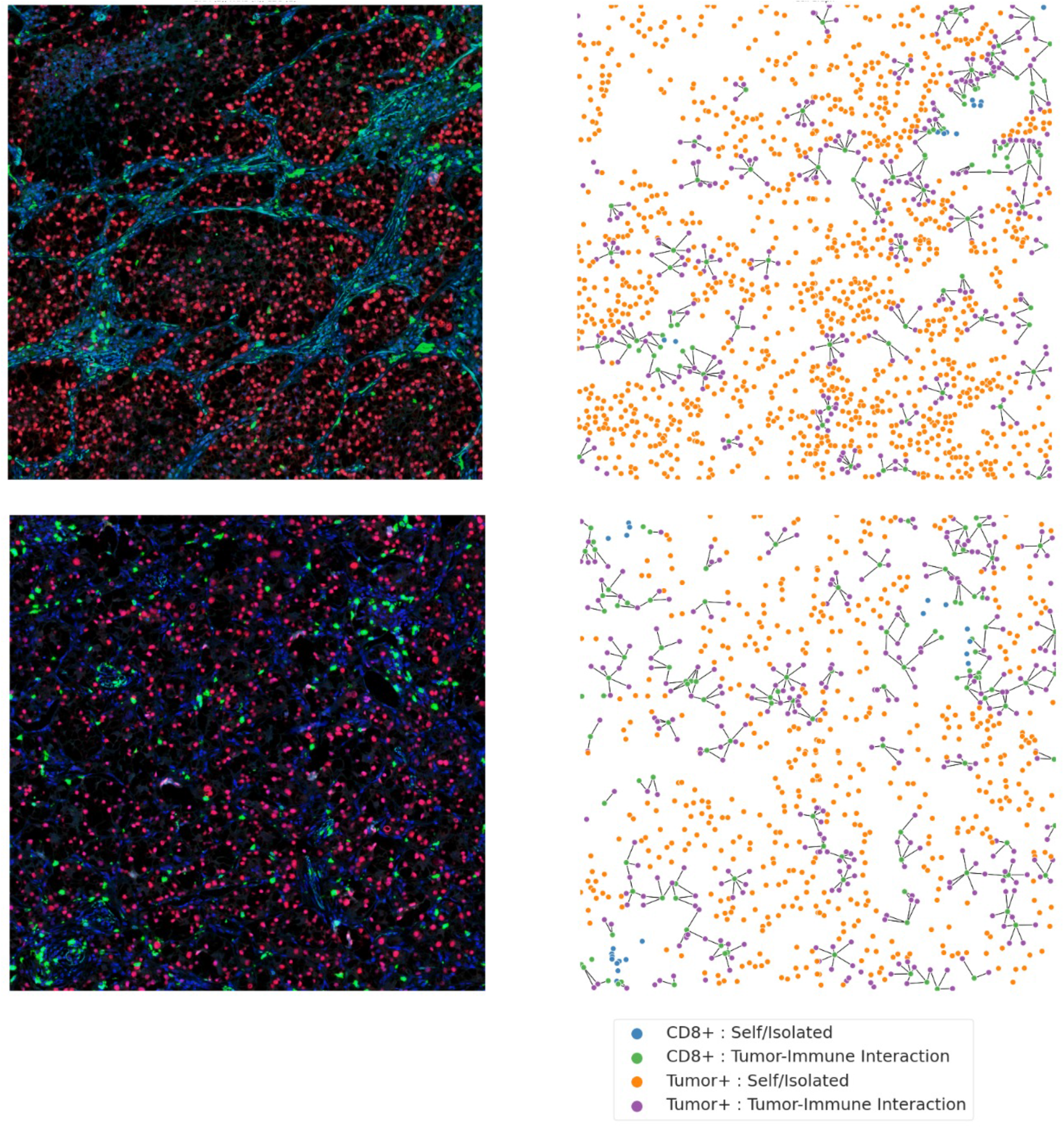
mIF data and cell graphs for immune hotspots: examples from two microhomogeneous cases. Edges are drawn between CD8+ and tumor cells that are adjacent in a nearest neighbor graph.

**Extended Data Figure 30:**
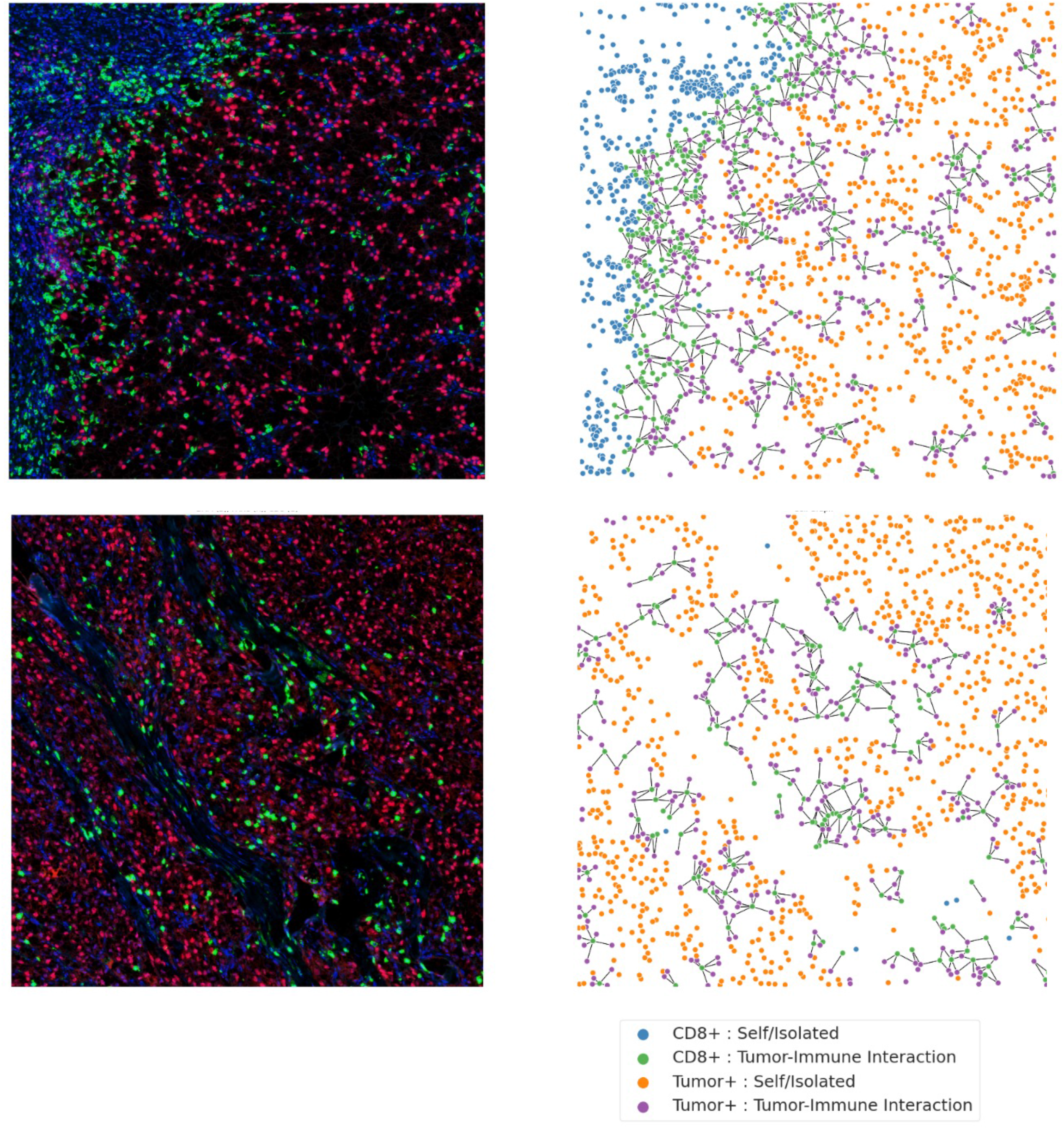
mIF data and cell graphs for immune hotspots: examples from two microheterogeneous cases. Edges are drawn between CD8+ and tumor cells that are adjacent in a nearest neighbor graph.

**Extended Data Fig. 31:**
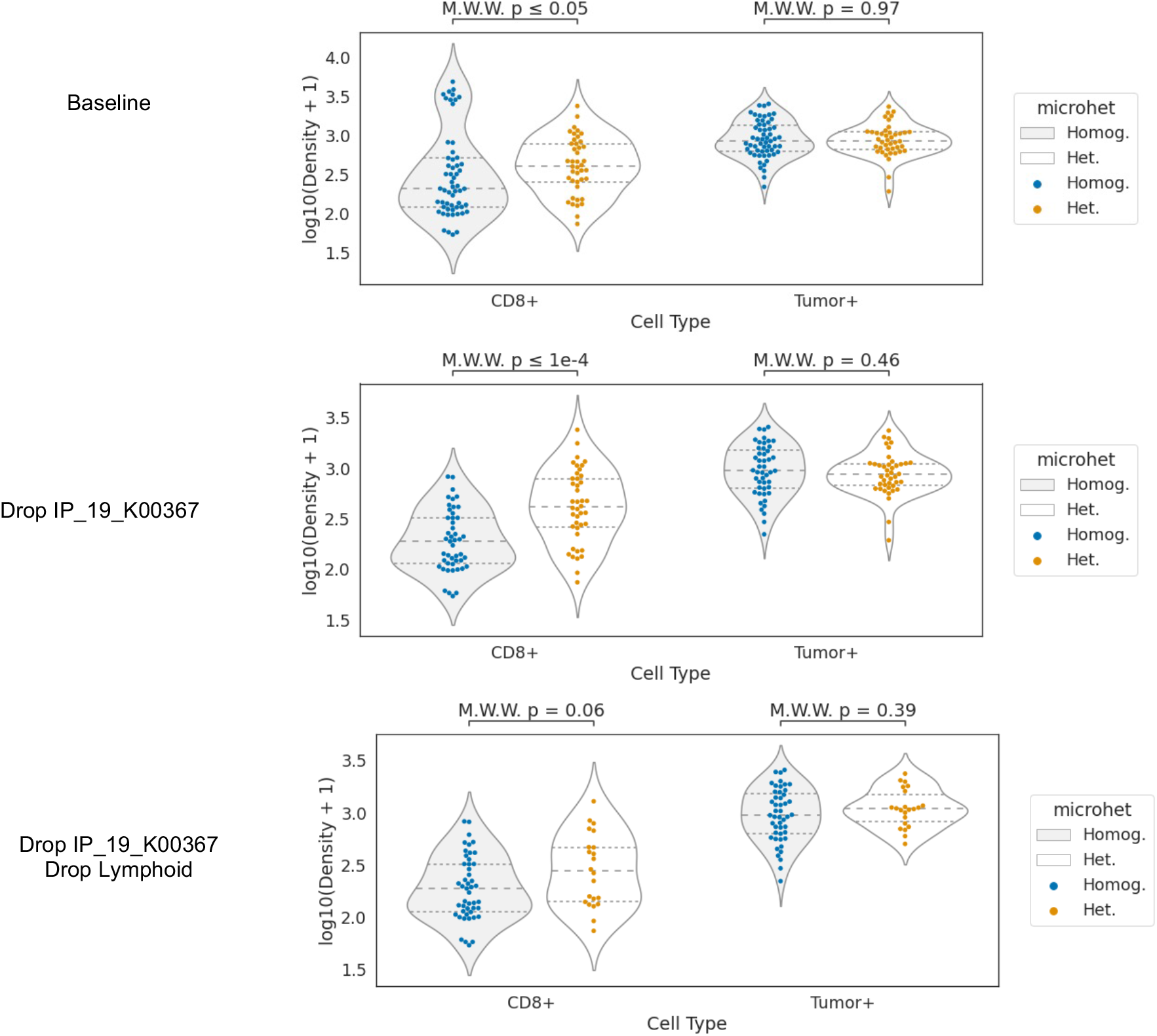
Cell densities by type and microheterogeneity status in immune hotspots. Y-axis: log10 density (cells per 2000px window; approx. 1mm) Rows: different data removal strategies. Significance calculated by Wilcoxon rank sum test (MWW).

**Extended Data Fig. 32:**
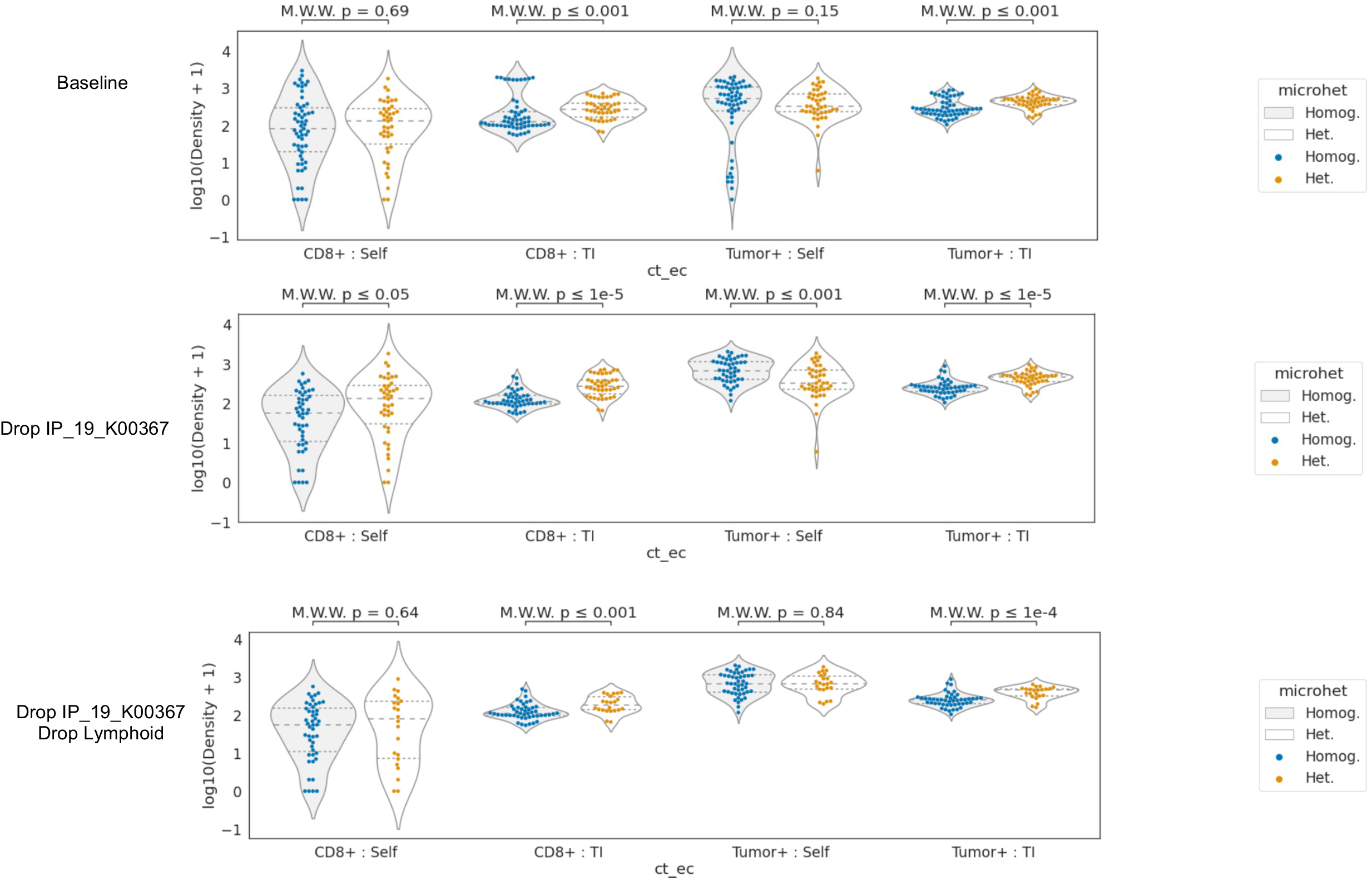
Cell densities by type, context, and microheterogeneity status in immune hotspots. Y-axis: log10 density (cells per 2000px window; approx. 1mm) Rows: different data removal strategies. Significance calculated by Wilcoxon rank sum test (MWW). “TI”: tumor-immune interacting cell context. “Self”: self-interacting cell context.

**Extended Data Fig. 33:**
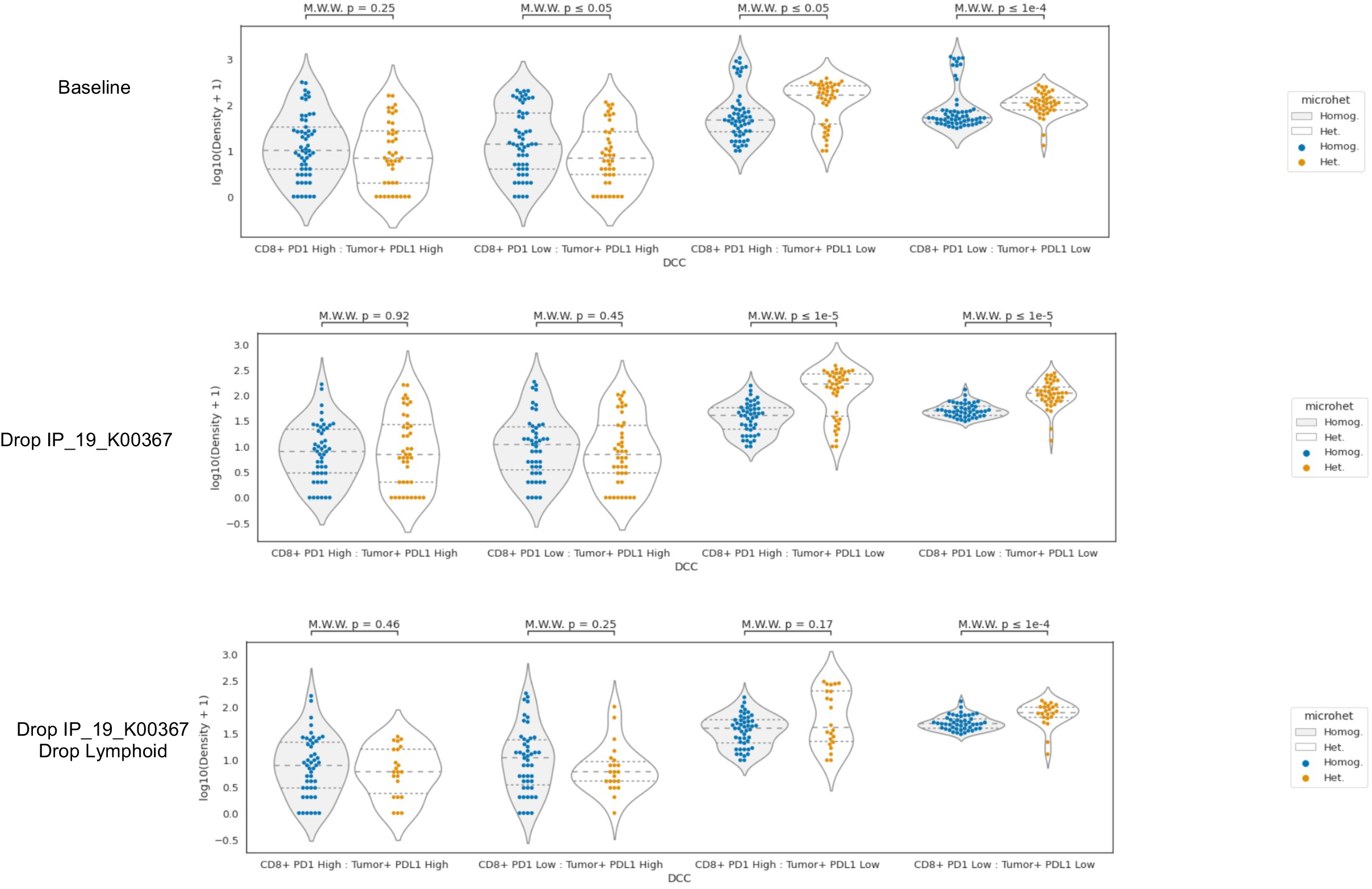
Cell densities by cell subtype and microheterogeneity status in immune hotspots. Y-axis: log10 density (cells per 2000px window; approx. 1mm) Rows: different data removal strategies. Significance calculated by Wilcoxon rank sum test (MWW).

